# Targeting CLK1 by CRISPR tools and pharmacological inhibition modulates the innate immune response and dengue virus replication

**DOI:** 10.64898/2026.04.15.718750

**Authors:** Berta Pozzi, Agustina Magalí Lucero, Anabella Srebrow

**Author notes:** Correspondence should be addressed to: Berta Pozzi,; Anabella Srebrow.

## Abstract

Alternative splicing is a key regulatory mechanism known to be altered upon viral infection. These alterations can arise both from direct viral interference with the splicing machinery and from cellular responses such as activation of innate immunity. Here, we investigated splicing changes shared between dengue virus (DENV) infection and interferon (IFN) treatment in cultured cells, the latter serving as a model for virus-independent innate immune activation. Among the common events, we identified an increased production of non-coding mRNA isoforms from the *CLK1* gene, which gives rise to a kinase that phosphorylates splicing factors including SR proteins and spliceosomal components. Consistent with this finding, IFN treatment led to a reduction in CLK1 protein levels. Using stable cell lines with CRISPR/dCas9-mediated modulation of *CLK1* expression, we found that silencing *CLK1* enhanced the induction of immune response genes, while its over-expression attenuated it. Inhibition of CLK1 kinase activity with the pharmacological inhibitor TG003 further potentiated IFN-induced gene expression and reduced DENV replication. Altogether, these results identify CLK1 as a proviral negative regulator of IFN-stimulated gene expression and suggest that its inhibition could enhance antiviral defenses and become a target for therapeutic strategies.

## Introduction

Dengue virus (DENV) is a growing threat to global health, with an estimated 400 million infections annually (1,2). Climate change is accelerating the spread of its mosquito vectors, such as *Aedes aegypti* and *Aedes albopictus*, leading to periodic outbreaks in various regions of the world (3). DENV is classified into four serotypes (DENV1-4), all with a single-stranded, positive-sense RNA genome of approximately 11 kb, which encodes a single polyprotein that is proteolytically processed into structural (C, PrM, E) and non-structural (NS1-NS5) proteins. The former constitute the viral capsid and envelope, while the latter perform diverse functions along the viral replication cycle (4,5). NS5 is the viral RNA polymerase and, although its function in viral genome replication is mostly cytoplasmic, it is predominantly located in the nucleus of cells infected with DENV2, compared to the other serotypes (6,7).

Our laboratory has demonstrated that DENV2 NS5 physically interacts with components of the cellular spliceosome, mainly with the CD2BP2 and DDX23 proteins of the U5 snRNP, interfering with the activity of this complex molecular machinery, consequently altering cellular splicing patterns and promoting a cellular environment more favorable for viral replication and proliferation (8). Nevertheless, altered alternative splicing (AS) in host cells is not exclusive to DENV but a common feature of many viral infections, as advances in RNA sequencing technologies have revealed for hundreds of host genes. These changes can arise from direct tampering of the splicing machinery by the virus, as with DENV NS5, or indirectly, as a consequence of activation of the innate immune response or cell damage induced during infection (9).

Activation of innate immunity requires the detection of viruses by cellular sensors and constitutes the first line of defense against these pathogens in human cells. In the case of RNA viruses, the detection of the viral genome in the cytoplasm is carried out by a group of receptors expressed by most cells of the body, known as RLRs (RIG-I-like receptors), including RIG-I (Retinoic Acid-Inducible gene I), MDA5 (Melanoma Differentiation-Associated protein 5), and LGP2 (Laboratory of Genetics and Physiology 2) (10). It was recently discovered that the interferon response induced by DENV infection depends on both RIG-I and MDA5 (11). Activation of RLRs by viral infection triggers signaling pathways that promote the expression of type I interferon (IFN-I), particularly IFNα and IFNβ, and other inflammatory cytokines. IFN secreted into the extracellular environment acts in an autocrine and paracrine manner, promoting the transcriptional increase of hundreds of IFN-inducible genes called “ISGs” (Interferon Stimulated Genes), through the INFAR1/2-JAK1-STAT1/2 pathway. Expression of ISGs leads to the establishment of an antiviral cellular environment that interferes with different stages of viral replication, restricting pathogenesis (12,13).

Here, we evaluated changes in splicing patterns shared between DENV infection and IFN treatment of cultured cells—the latter being a way to activate the innate immune response in the absence of viral components. Among the shared events, we identified an increase in non-coding mRNA isoforms derived from the *CLK1* gene, which encodes for a kinase that phosphorylates splicing factors, including SR proteins and the spliceosomal protein U1-70K. These proteins are involved in the regulation of the splicing process, and their phosphorylation by CLK1 affects their recruitment to nuclear speckles, dynamic reservoirs of splicing factors that modulate mRNA isoform production and consequently protein diversity (14).

After confirming a reduction in CLK1 protein levels following interferon treatment of cultured cells, we set out to analyze the induction of immune response genes in stable cell lines in which CLK1 levels were modulated using the dCas9-KRAB/VP64 system. We observed that the induction of immune response genes increased upon CLK1 silencing, whereas the opposite effect was observed with CLK1 over-expression. We also analyzed the induction of these genes in the presence of TG003, an inhibitor of CLK1 kinase activity, and found that interferon-induced expression was further enhanced. Furthermore, we measured viral replication in TG003-treated, dengue-infected cells. We observed a significant reduction in viral load, confirming that modulation of interferon response genes by CLK1 exerts an antiviral effect.

## Materials and methods

### RNA-seq data acquisition and analysis

RNA-seq .fastq files were obtained from Gene Expression Omnibus (GEO) for the sequencing of DENV-infected or mock-infected A549 cells at 24 hours post-infection (GEO access: GSE84285) (8), and from Peripheral Blood Mononuclear Cells (PBMCs) treated with IFN or untreated (GEO access: GSE72502) (15). The workflow depicted in Figure 1a was followed, using Galaxy, a free web platform designed to facilitate bioinformatics analysis, and subsequently the 3D-RNA-seq tool (16). Genes with Differential Alternative Splicing (DAS) upon DENV infection and IFN treatment are listed in Tables S1 and S2.

**Figure 1:**
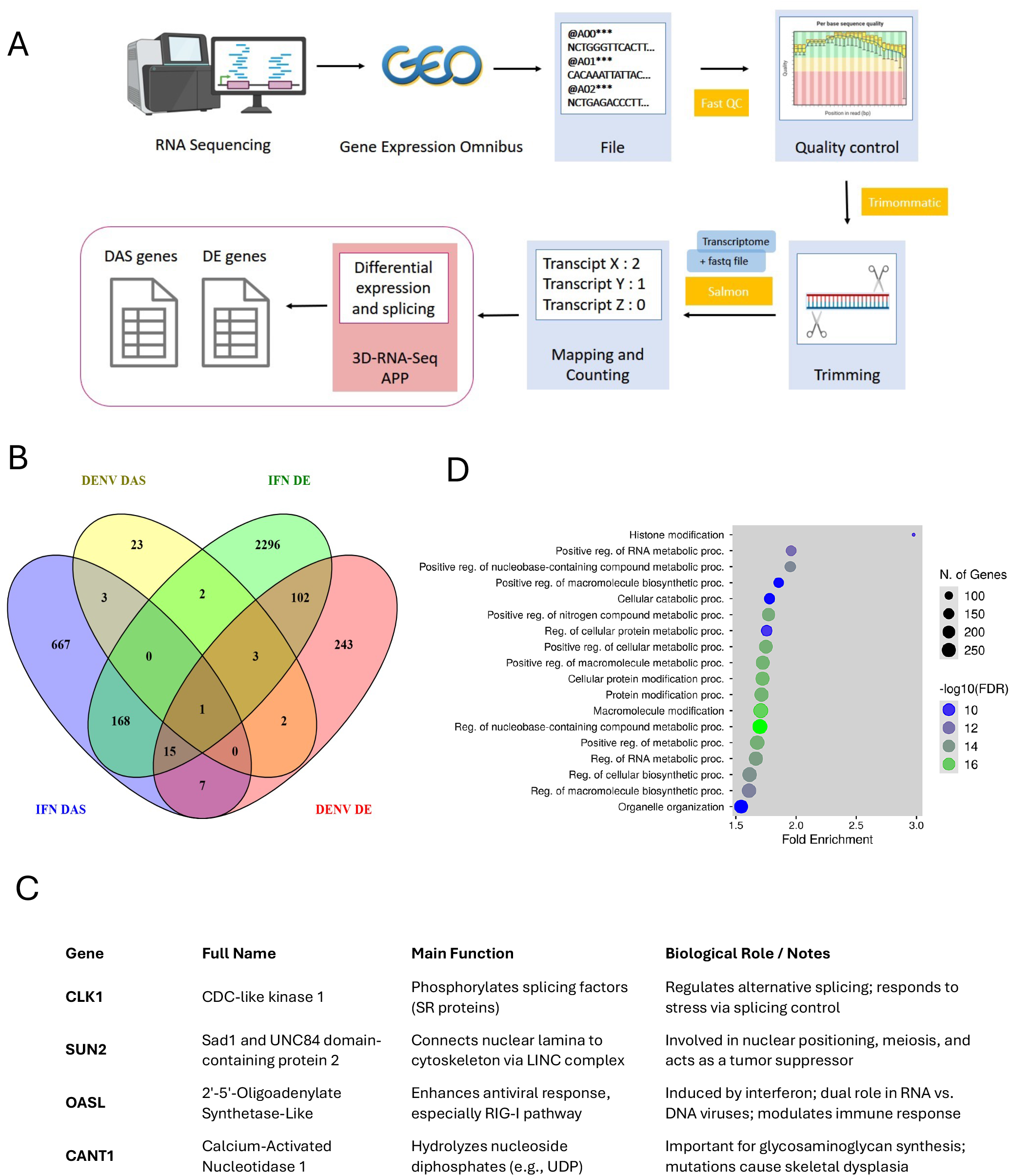
Genes with Differential Expression (DE) and Differential Alternative Splicing (DAS) upon DENV infection and IFN treatment. A) Pipeline for analysis of publicly available databases GSE84285 (A549 cells RNA-seq infected with Dengue virus 2 or mock for 24 hours) and GSE72502 (Healthy donor PBMC RNA-seq with or without IFN-alpha stimulation). B) Venn diagram for genes with DE and DAS upon DENV infection and IFN treatment. C) Biological functions of genes shared between both. D) Gene Ontology analysis of DAS genes upon IFN treatment.

### Functional category analysis (Gene Ontology)

The list of genes with Differential Alternative Splicing (DAS) upon IFN treatment was subjected to a functional category analysis, or Gene Ontology analysis; using “ShinyGO v0.741: Gene Ontology Enrichment Analysis” (17).

### Cell culture and transfection

All cell lines used (HEK293T derived from human embryonic kidney cells and A549 derived from lung adenocarcinoma) were maintained in DMEM medium, supplemented with 10% fetal bovine serum, 4.5 g/L glucose, 110 mg/L sodium pyruvate, 100 U/mL penicillin, and 100 μg/mL streptomycin, in an incubator at 37°C with 5% CO2. HEK293T cells were transfected with polyethyleneimine (PEI), according to manufacturer’s instructions (Polyciences).

### DENV infections and treatment with IFN and TG003

DENV2 infections, strain 16681, were performed with a multiplicity of infection (MOI) of approximately 1. Viral adsorption was carried out at 4°C with shaking, and after one hour of incubation, the medium was replaced with fresh medium. For IFN treatments, 50,000 U/ml of interferon α2b (Biosidus) was added to the cell culture medium. Treatment with the CLK1 inhibitor (TG003) was performed at a final concentration of 1 μM for 72 h, and the corresponding control was performed with the appropriate volume of DMSO.

### RNA extraction, reverse transcription and qPCR

RNA was extracted using TRIpure reagent (Roche), following the manufacturer’s protocols. To obtain cDNA from total RNA, the MMLV reverse transcriptase enzyme (Inbio Highway) was used, following the manufacturer’s protocol. For qPCRs, 2.5 μl of a 1:20 dilution of the RT reaction containing cDNA was used as template combined with 10 μl of a mixture with: PCR Buffer 1.25 μl, MgCl_2_ 25 mM 2 μl, Primer Fw 10 μM 0.5 μl, Primer Rev 10 μM 0.5 μl, dNTPs 10 mM 0.25 μl, Taq DNA polymerase (5 U/μl) 0.05 μl, Sybr Green 0.0125 μl and H2O 5.94 μl, in a Mastercycler® ep Realplex PCR device (Eppendorf). The annealing temperature was 60°C and the elongation time at 72°C was 30 s. Relative RNA abundance from cDNA samples and no-reverse transcription controls was estimated employing internal standard curves with a PCR efficiency of 100 ± 10% for each set of primers, in each experiment. Realplex qPCR software was used to analyze the data. All primers used are listed in Table S3.

### Western Blot

Protein samples were resolved by SDS-PAGE and transferred to nitrocellulose membranes (BioRad). Membranes were blocked and then incubated with primary antibodies. After washing, membranes were incubated with IRDye® 800CW or 680RD (LI-COR Biosciences) secondary antibodies. Bound antibody was detected using an Odyssey imaging system (LI-COR Biosciences). Western blots were performed at least three times from independent experiments and representative images are shown in each case. The primary antibodies used were: Anti-CLK1 antibody (ab74044), anti-HA (Covance) and mouse monoclonal anti-vinculin hVIN-1 (Sigma-Aldrich).

### DNA plasmid expression vectors

HA tagged CLK1 was obtained by cloning CLK1 cDNA into pcDNA/FRT with an HA coding forward primer. The reporter constructs were obtained by replacing the SFFV promoter of PHR-SFFV with a 400 bp region of the IFIT1 promoter upstream of GFP coding sequence. ISG15 intron was cloned downstream of GFP using a BamHI site introduced by PCR. All primers are listed in Table S3.

### CRISPRa and CRISPRi systems

The CRISPRa system was used for activation and CRISPRi for inhibition of CLK1 transcription. First, stable cell lines expressing dCas9-KRAB and dCas9-VP64 were obtained by lentiviral transduction and cytometric selection, taking into account that these vectors express fluorescent markers. The Benchling platform was used to design the single guide RNAs (sgRNAs) (Benchling, 2025), within a 300 bp region of the CLK1 gene promoter (18) (Table S3). An sgRNA with no homology in the genome (sgNT, *Non-Target*) was used as control. The candidate guides were cloned into the pLentiGuide Puro vector (Addgene #52963) and co-transfected with the dCas9-KRAB/dCas9-VP64 vectors (Addgene #60954 and #61422) in HEK293T. CLK1 silencing/over-expression was evaluated by RT-qPCR and the best sgRNA was selected to generate stable cell lines by transduction of previously generated dCas9-KRAB/dCas9-VP64 lines and selection with puromycin. In all cases, to obtain the lentiviruses, the lentiviral vectors were co-transfected with packaging, envelope, and regulatory vectors using PEI in HEK293T cells.

### Flow cytometry

HEK293T cells transfected with the fluorescent reporter were harvested in phosphate-buffered saline (PBS) to a total volume of 500 µL to obtain 1 × 10^6^ cells mL^−1^. Data were acquired using a BD FACSAria II Flow Cytometer (Becton Dickinson) and analyzed using BD FACSDiva and FlowJo v10 softwares.

### Statistics

Typically, three independent experiments with triplicates were performed for each condition evaluated. Mean values with their standard deviations and P-values, calculated using a paired two-tailed t-test, are presented. Significant P-values are indicated with asterisks on the graphs (***P < 0.001; **P < 0.01; *P < 0.05).

## Results

### CLK1 splicing patterns change with viral infection and the innate immune response

We set out to identify splicing changes in host genes upon viral infection and induction of the innate immune response. RNA-seq data from A549 cells infected with DENV or mock-infected were analyzed 24 hours post-infection (GEO accession: GSE84285), as well as data from PBMCs treated with IFN or untreated (GEO accession: GSE72502). Analysis of these datasets, following the pipeline depicted in Figure 1a, revealed genes with differential expression (DE) in response to DENV infection (DENV DE) and interferon treatment (IFN DE), as well as genes with differential alternative splicing (DAS) in both contexts (DENV DAS and IFN DAS) (Table S1 and S2). To explore the overlap between these conditions, a Venn diagram was constructed (Fig. 1a), revealing that 102 genes exhibit differential expression in both IFN-treated and DENV-infected cells, while only 3 genes share differential alternative splicing events in both conditions (Fig. 1b). The latter correspond to CLK1, SUN2, and CANT1 (Fig. 1c).

On the other hand, a single gene was detected that simultaneously exhibits differential expression and differential alternative splicing under both treatments. This gene corresponds to OASL, a member of the 2′-5′ oligoadenylate synthase (OAS) family, known for its antiviral role. OASL regulates the early phase of infection by degrading viral RNA in conjunction with RNase L, thereby blocking viral replication (19). The finding of OASL as a gene with regulation at both the DE and DAS levels in both conditions indicates that the approach used is effective in detecting genes with functional antiviral potential.

Using a gene ontology analysis (ShinyGO 0.85.1), the biological functions enriched in DAS genes by IFN treatment were explored (Fig. 1d). We observed functional categories related to transcription (such as histone modifications or RNA metabolism) but also positive regulations of metabolic processes and protein modifications. Remarkably, CLK1 was included in several of these categories given its widespread role as a kinase linking extracellular stimuli with the RNA processing machinery. Indeed, CLK1 activity was reported to be regulated by body-temperature (20) and to be responsible for temperature-dependent modulation of splicing patterns . Furthermore, previous findings have shown that its silencing affects the replication of certain viruses, such as influenza A, reinforcing its potential antiviral relevance (21).

### Alternative splicing of CLK1 pre-mRNA in response to DENV and IFN

The *CLK1* gene (ENSG00000013441) gives rise to a coding mRNA isoform where all annotated exons are preserved, whereas non-coding isoforms arise either by retention of introns 3 and 4 or exclusion of exon 4. In the later one, a premature STOP codon renders this isoform for degradation by Non-sense Mediated mRNA Decay (NMD) (Fig. 2a). CLK1 exhibits differential alternative splicing (DAS) in response to both DENV infection and IFN treatment. Analysis of the relative abundance of CLK1 isoforms using TPM values (Transcripts Per Million) revealed that the coding isoform (indicated in red) decreased in both treatments, while the isoforms associated with intron retention (IR) and exon skipping (ES) increased (Fig. 2b and c). This pattern suggests a possible regulatory mechanism for CLK1 under antiviral stress conditions. A similar finding was described by Uzor et al., 2018 (22), who observed negative autoregulation of CLK1 in cells exposed to various types of environmental stress.

**Figure 2:**
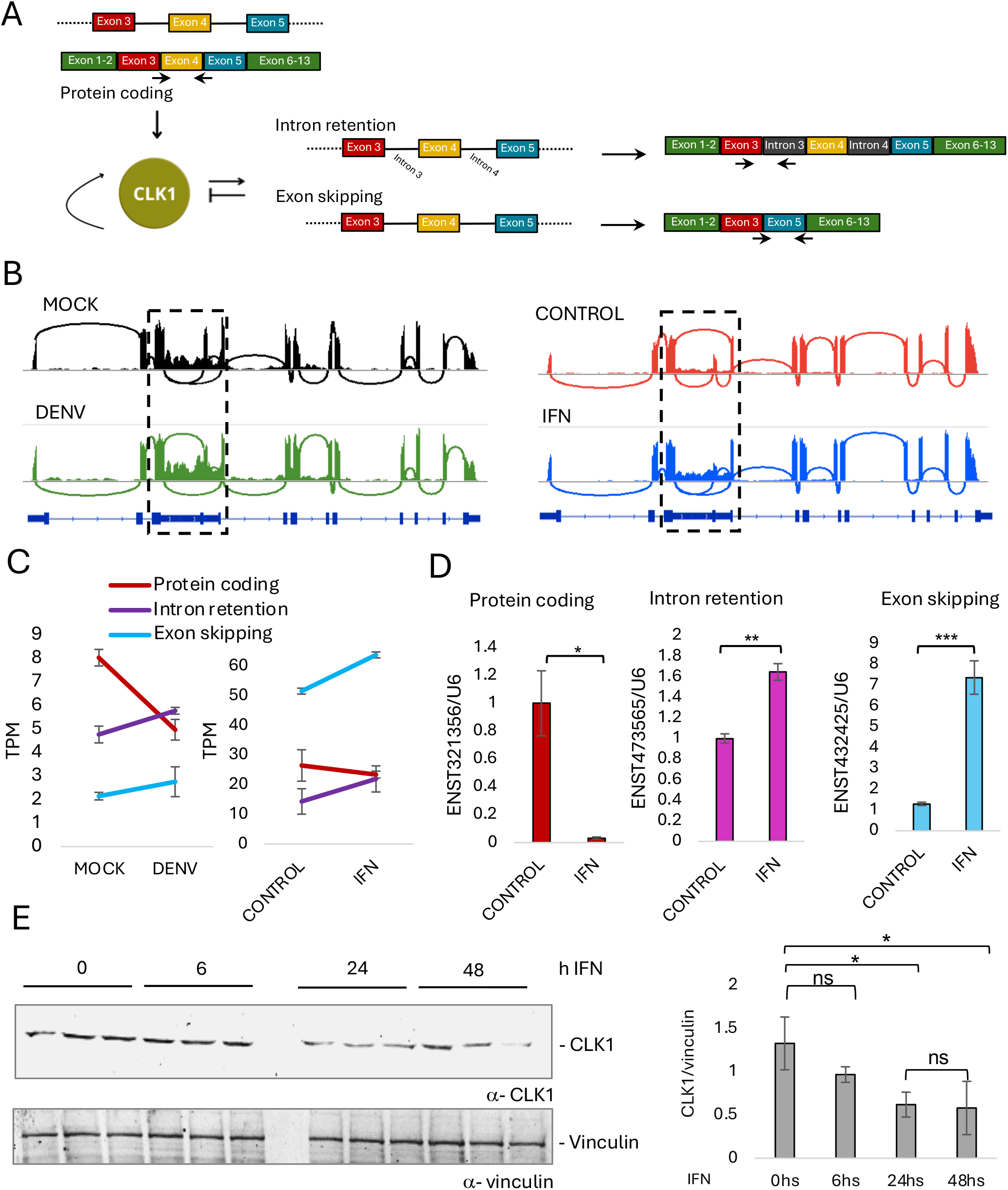
CLK1 pre-mRNA alternative splicing is modulated upon DENV infection and IFN treatment. A) Model depicting CLK1 pre-mRNA alternative splicing isoforms generated by intron retention and exon skipping. B) Sashimi plots of CLK1 RNA-seq read coverage, showing intron retention and exon 4 skipping upon DENV infection and IFN treatment (dashed box). C) Quantification in Transcripts Per Million (TPM) of CLK1 annotated isoforms with retained introns and skipped exon 4. D) Validation of isoform abundance by RT-qPCR with specific primers spanning the different isoforms, indicated in the scheme in A, upon IFN alpha treatment (50000UI/ml) of A549 cells. E) Determination of CLK1 protein levels by western blot. Significant P-values, calculated using a paired two-tailed t-test, are indicated with asterisks on the graphs (***P < 0.001; **P < 0.01; *P < 0.05).

RNA-seq results were validated with specific primers for all CLK1 isoforms by RT-qPCR upon treatment with recombinant IFN alpha (50,000 U/ml for 24 h) in cultured A549 cells. Similar patterns were observed: a decrease in the coding isoform and an increase in isoforms with intron retention or exon skipping (Fig. 2d).

With the aim of analyzing whether the decrease of the coding mRNA isoform has an impact on CLK1 protein levels, a western blot was performed at different time points (0, 6, 24 and 48) of IFN addition (50,000 U/ml) in A549 cultured cells (Fig. 2e). Quantification showed that after 6 hours of IFN treatment, CLK1 abundance was comparable to the control, but a significant decrease was observed at 24 and 48 hours. Furthermore, no significant changes in CLK1 protein levels were noted between 24 and 48 hours; therefore, it was decided to continue with 24-hour treatments from this point onward.

### CLK1 silencing enhances ISG expression

In an attempt to knock-down CLK1, we employed the CRISPRi system. We evaluated the efficiency of five different sgRNAs, designated sg1 to sg5, by transiently co-transfecting them into HEK293T cells with a vector encoding for dCas9-KRAB (Fig. 3a). At 48 hours post-transfection, total RNA was extracted, and CLK1 mRNA levels were quantified by RT-qPCR. As shown in Figure 3b, sg4 and sg5 induced a significant decrease in CLK1 coding isoform expression compared to the control sgRNA (sgNT, for Non-Target single guide), which has no complementary sequences in the human genome. Based on this proof of concept, we generated a stable cell line by lentiviral transduction of sg4 into A549 cells stably expressing dCas9-KRAB. As a control, a cell line transduced with the sgNT was obtained (Fig 3c).

**Figure 3:**
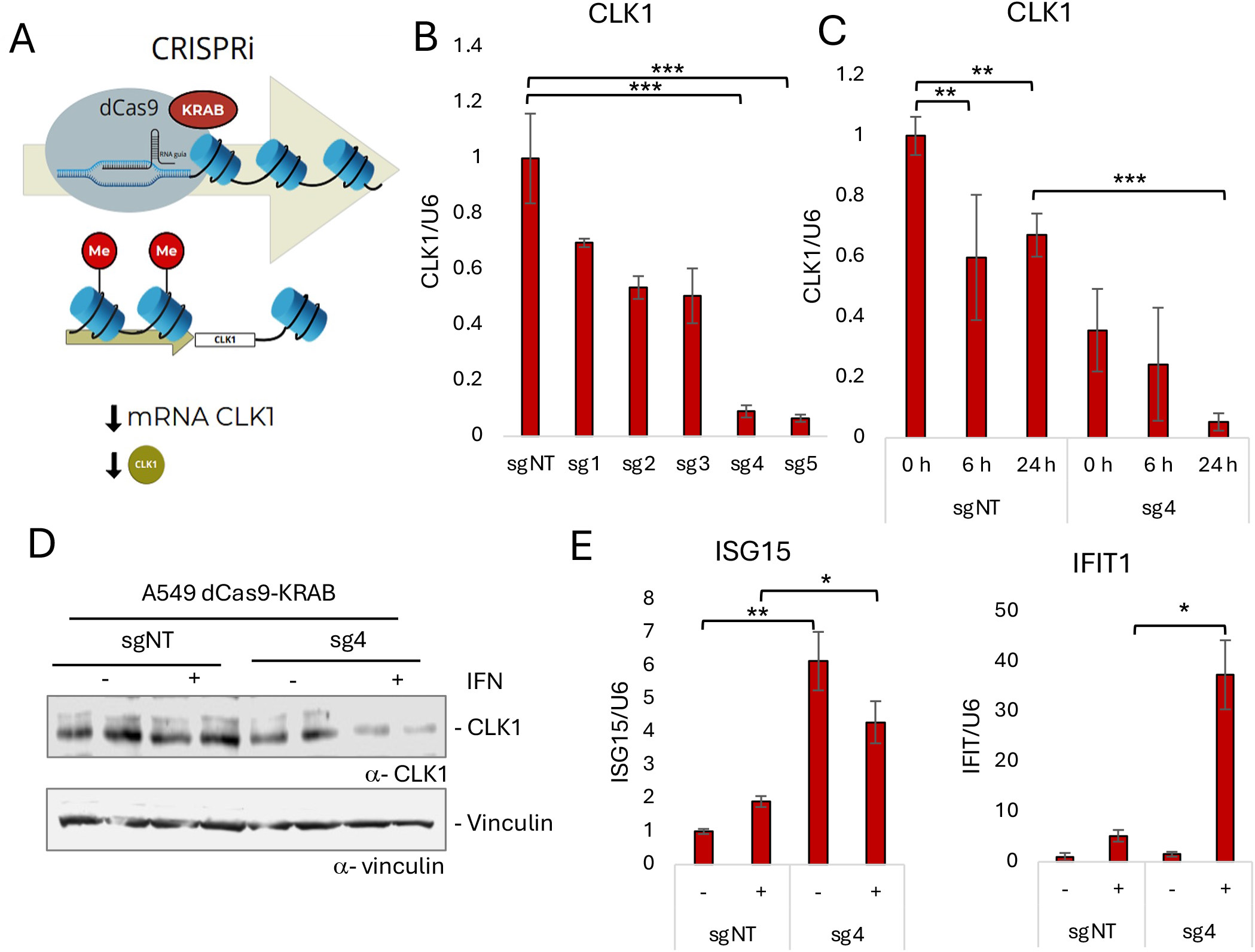
Interferon Stimulated Genes expression is enhanced by CLK1 CRISPRi. A) Model depicting CRISPRi design for CLK1 silencing. B) Quantification of CLK1 coding isoform by RT-qPCR upon transient transfection of five different sgRNAs spanning CLK1 proximal promoter region, in comparison with an sgRNA with no homology in the genome (sgNT, Non-Target) in HEK293T cells. C) CLK1 mRNA quantification by RT-qPCR in a stable A549 cell line transduced with dCas9-KRAB and sgRNA 4 or NT and treated with recombinant IFN alpha for the indicated hours. D) CLK1 protein levels by western blot upon CLK1 CRISPRi and 24 h IFN alpha treatment. E) Quantification of ISG15 and IFIT1 upon CLK1 CRISPRi and 24 h IFN alpha treatment by RT-qPCR. Significant P-values, calculated using a paired two-tailed t-test, are indicated with asterisks on the graphs (***P < 0.001; **P < 0.01; *P < 0.05).

To evaluate the combined effect of CRISPRi mediated silencing and IFN treatment on CLK1 expression, 50,000 U/ml of IFN was added to the culture media of the former generated stable A549 cell lines. After 24 hours of incubation, total RNA was extracted, reverse transcribed, and CLK1 mRNA levels were quantified by RT-qPCR (Figure 3c). As previously observed, IFN exposure resulted in decreased levels of CLK1 mRNA (coding isoform) at both 6 and 24 hours of treatment. Furthermore, CRISPRi significantly reduced CLK1 expression when comparing sg4 to sgNT, demonstrating a specific silencing effect, which was even more pronounced at 24 hours of IFN treatment. This indicates an additive effect between IFN treatment and CRISPRi on CLK1 mRNA expression levels. To verify that the system had silenced CLK1 not only at the mRNA level but also at the protein level, a western blot was performed using whole-cell lysates from A549 stably expressing dCas9-KRAB and either sgNT or sg4 and treated with IFN for 24 hours. The additive effect of CRISPRi and IFN was also observed at protein level, using vinculin as a loading control (Fig. 3d).

The host response to a viral infection includes the induction of IFN-I and the activation of hundreds of ISGs as part of the antiviral program, as mentioned previously. Among them, ISG15 encodes a ubiquitin-like protein, whose expression is strongly induced by IFN and which has been implicated as a central player in antiviral defense (23). IFIT1 represents another classic ISG, highly sensitive to type I IFN stimulation. IFIT1 transcription is rapidly induced and reaches high levels in many cell types after viral infection or exogenous IFN administration, and is therefore frequently used as a marker of IFN pathway activation and viral infection (24).

Therefore, to evaluate the effect of CLK1 silencing and IFN stimulation on the expression of ISGs, cell lines stably expressing dcas9-KRAB and either sg4 or sgNT were used. After 24 hours of IFN treatment, ISG15 and IFIT1 mRNA levels were quantified by RT-qPCR (Figure 3e). We observed that, as expected, IFN treatment leads to an increase in the expression of both ISGs. Furthermore, we observed an enhanced induction upon CLK1 silencing by CRISPRi. These results indicate that modulating CLK1 expression levels has an impact on the induction of immune response genes.

### CLK1 over-expression attenuates ISG expression

The CRISPRa system (25) was used to over-express CLK1 (Fig. 4a) by generating a stable A549 cell line for the fusion protein dCas9-VP64 and sg4 or the NT guide. An increase in CLK1 mRNA levels (coding isoform) was observed, when comparing sg4 containing cells with the sgNT ones (Fig. 4b). This is consistent with the protein levels observed by western blot, where also over-expression of CLK1 is attenuated when combined with IFN treatment (Fig 4c).

**Figure 4:**
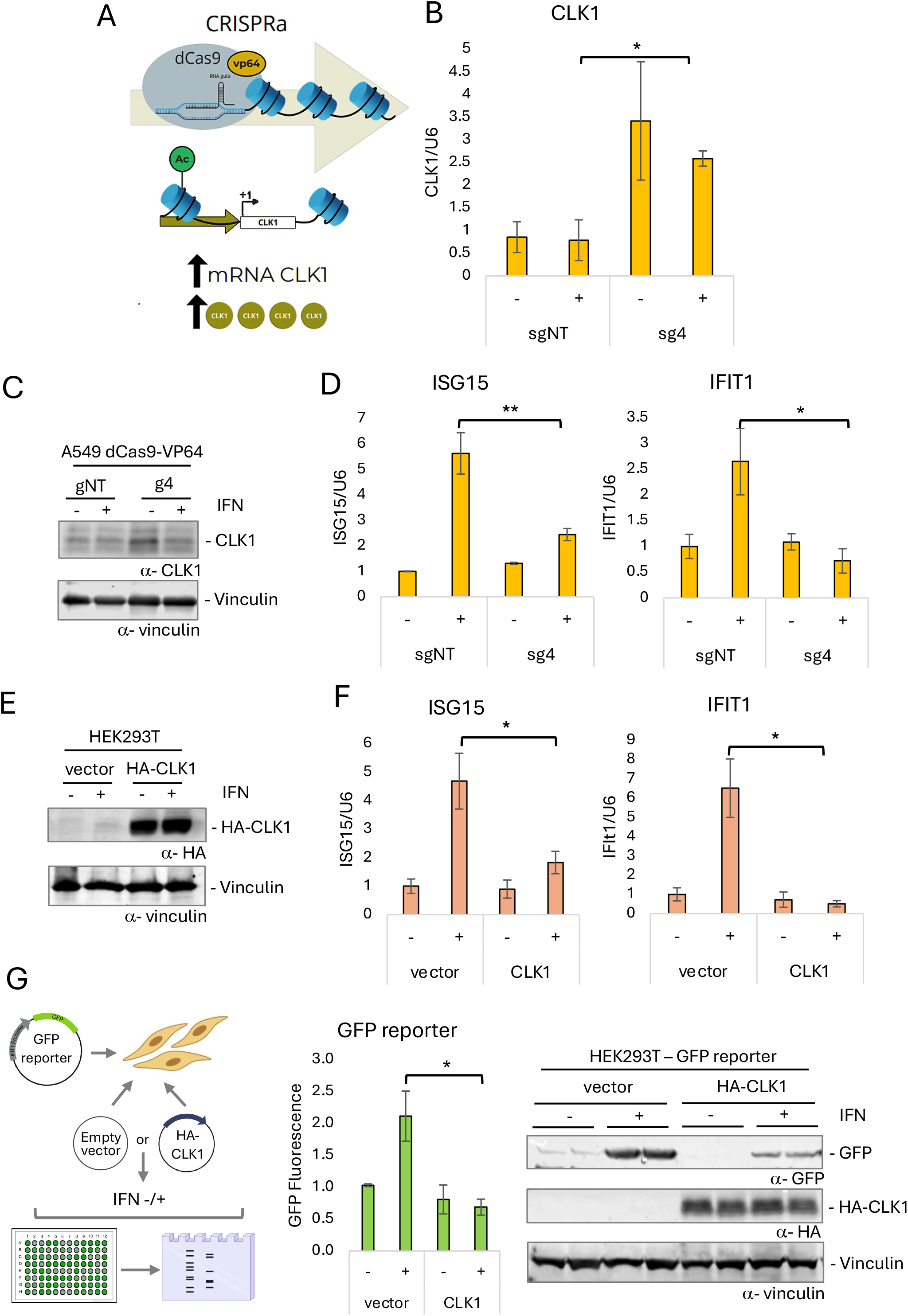
Interferon Stimulated Genes expression is attenuated by CLK1 CRISPRa. A) Model depicting CRISPRa design for CLK1 over-expression. B) CLK1 mRNA quantification by RT-qPCR in a stable A549 cell line transduced with dCas9-VP64 and sgRNA 4 or NT and treated with recombinant IFN alpha for the indicated hours. C) CLK1 protein levels by western blot upon CLK1 CRISPRa and 24 h IFN alpha treatment. D) Quantification of ISG15 and IFIT1 upon CLK1 CRISPRa and a 24 h IFN alpha treatment by Rt-qPCR. E) CLK1 protein levels by western blot upon transient transfection with empty or HA-CLK1 coding vectors and 24 h IFN alpha treatment. F) Quantification of ISG15 and IFIT1 by RT-qPCR upon HA-CLK1 over-expression and 24 h IFN alpha treatment in HEK293T cells. G) Determination of GFP expression driven by IFIT1 promoter by fluorescence microplate reader and western blot, with two replicates for each experimental condition. The reporter was transiently transfected in HEK293T cells together with empty vector or HA-CLK1 for 48 h and treated with IFN for the last 24 h. Significant P-values, calculated using a paired two-tailed t-test, are indicated with asterisks on the graphs (***P < 0.001; **P < 0.01; *P < 0.05).

Given that the CRISPRa cell line targeting CLK1 exhibited a slow growth phenotype, we decided to test over-expression by transient transfection of a CLK1 expression vector into HEK293T cells. A western blot was performed under IFN-treated conditions and using an empty vector as a control. As shown in Figure 4d, the transfection of the plasmid increased CLK1 protein levels, as expected. Analogous to the silencing assays, the expression levels of the same ISGs (ISG15 and IFIT1) were evaluated in response to CLK1 over-expression, either using the CRISPRa system or the over-expression vector. As shown in Figure 4e, while IFN treatment increased the expression of both genes, CLK1 over-expression significantly attenuated this induction. Consistently, the same pattern was observed when over-expression was achieved through transient transfection with a CLK1 expression vector (Fig. 4f). Furthermore, this HA-tagged vector was used for reporter assays, by being co-transfected with a plasmid where the expression of GFP is driven by the IFIT1 promoter. As shown in Fig 4g, the expression levels of GFP upon IFN treatment decrease when CLK1 is over-expressed, again indicating that CLK1 attenuates the induction of ISGs. This result was corroborated by measuring GFP fluorescence in live cells with a microplate reader and by western blot with an anti-GFP antibody (Fig 4g).

### TG003 enhances innate immune response and attenuates viral replication

For the purpose of modulating CLK1 activity, we used a commercial kinase inhibitor, known as TG003. Using cultured A549 cells, we performed a 24-hour treatment with TG003 followed by 24-h IFN incubation and assessed the induction of ISG15 and IFIT1. The combined treatment with IFN and the kinase inhibitor resulted in a marked increase in the expression of both selected ISGs (Fig 5a). This was also evaluated with the GFP reporter, noticing that GFP induction with IFN was higher when combined with TG003 (Fig 5b). This was also observed inducing the reporter together with the presence of a mutant version of CLK1, where 3 of its phospho residues (S36, T122 and S139) were replaced by alanine (Fig 5c). On the contrary, the phosphomimetic mutant abolished GFP expression. Altogether, these results suggest that CLK1 activity modulates the innate immune response.

**Figure 5:**
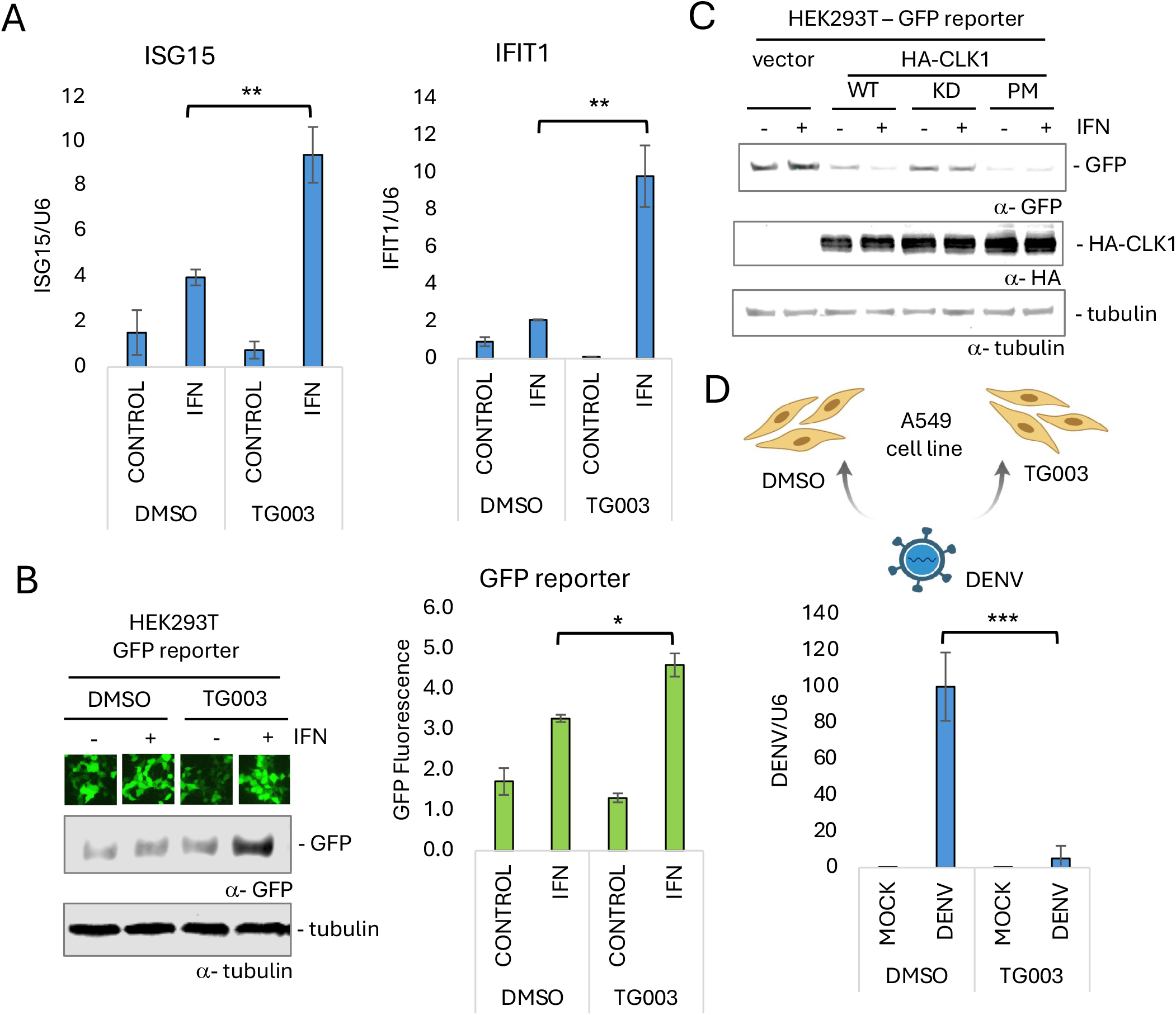
Pharmacological inhibition of CLK1 induces ISG expression and attenuates flaviviral infection. A) Quantification of ISG15 and IFIT1 by RT-qPCR upon treatment with TG003 (1 µM for 48 h) and IFN alpha (50000 UI/ml for 24 h). B) Determination of GFP expression driven by IFIT1 promoter by western blot and fluorescence microplate reader upon treatment with TG003 (1 µM for 48 h) and IFN alpha (50000 UI/ml for 24 h). C) Determination of GFP expression driven by IFIT1 promoter by western blot in the presence of HA-CLK1 wt or mutant versions and IFN treatment (50000 UI/ml for 24 h). D) Model depicting CLK1 pharmacological inhibition with TG003 (1 µM for 48 h) followed by DENV infection (48 hpi) and quantification of DENV RNA by RT-qPCR. Significant P-values, calculated using a paired two-tailed t-test, are indicated with asterisks on the graphs (***P < 0.001; **P < 0.01; *P < 0.05).

Finally, the impact of inhibiting CLK1 kinase activity on DENV-2 replication was assessed. A549 cells were treated with TG003 for 72 hours and infected with DENV-2 during the last 48 hours (Fig. 5d). Remarkably, the treatment with TG003 resulted in a significant reduction of viral load. This observation aligns with an emerging strategy targeting cellular kinases as antiviral therapeutic targets based on numerous studies that have demonstrated that viruses depend on several host kinases (such as CDK, MAPK, Src, JAK, and CK) for various steps in their multiplication cycle, including replication, assembly, and viral release (26). While further studies are needed to precisely elucidate the role of the CLK1 in DENV-2 replication cycle, our results suggest that the use of TG003 inhibitor represents a promising antiviral strategy.

### TG003 modulates ISG intron splicing

Considering that CLK1 modulates a plethora of SR protein phosphorylation and, consequently, of splicing events, it is tempting to speculate that CLK1 could modulate IFIT1 and ISG15 pre-mRNA splicing. Curiously, we investigated their exonic and intronic architecture to find out that the most IFN-induced transcripts of both ISGs contain a unique intron and that available RNA-seq data of cells treated with TG003 show less intronic reads of ISG15 upon treatment with this inhibitor (Fig. 6a) (20).

**Figure 6:**
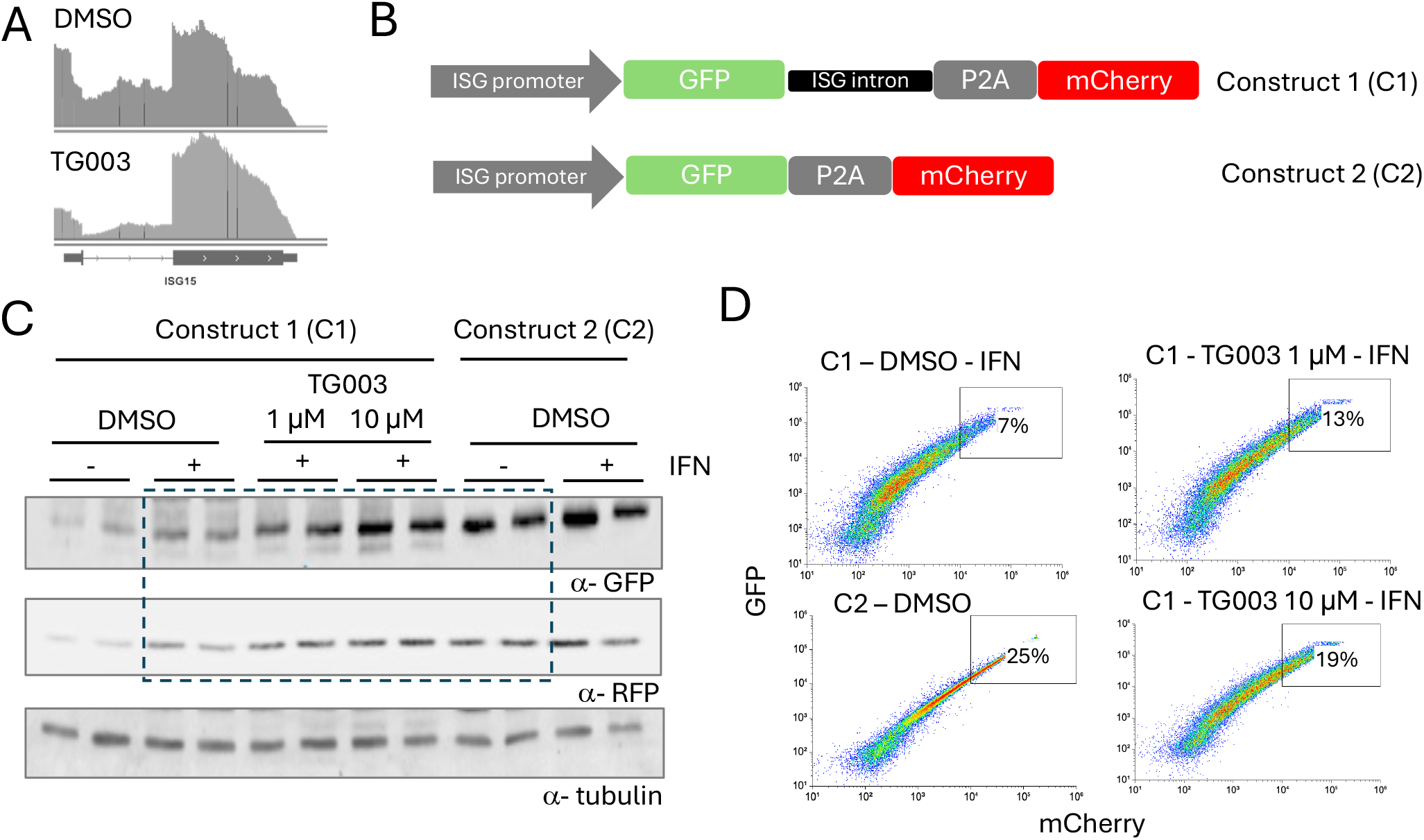
TG003 modulates ISG intron splicing. A) ISG15 RNA-seq read coverage, showing less intron reads upon TG003 treatment. B) Scheme of fluorescent reporters with ISG promoter (Construct 1 and 2) and intron (Construct 1). C and D) Determination of GFP and mCherry expression driven by constructs 1 and 2 by western blot (A) and flow cytometry (D) in the presence of TG003 (1 and 10 µM for 48 h) and IFN treatment (50000 UI/ml for 24 h). Flow cytometry charts belong to western blot samples depicted by the dashed box, with two replicates for each experimental condition.

Taking all these into account, we designed an intron-containing fluorescent reporter as depicted in Figure 6b, where expression is again driven by IFIT1 promoter but GFP and mCherry cassettes are separated by ISG15 intron and a P2A peptide (Construct 1) or the later alone (Construct 2). We transfected these constructs into HEK293T cells and evaluated GFP and mCherry expression levels by western blot (Fig. 6c) and flow cytometry (Fig. 6d), in the presence of IFN and TG003, as shown in each panel. When employing construct 1, containing the intron, both fluorescent markers were consistently induced by IFN, but this induction was stimulated even further (particularly for GFP) when IFN was combined with TG003, in a concentration dependent manner for the latter. Even more, induction levels at the higher dose of TG003 resembled that of construct 2, lacking the intron (Fig. 6c). These results evidenced that intron splicing is relevant for ISG expression and that its modulation can boost immune response. Furthermore, flow cytometry data suggest that intronic presence gives rise to a more variable GFP expression within the cell population (C1-DMSO, induced with IFN). When TG003 is added, and again in a dose dependent manner (C1-TG003 1 and 10 µM), this variability decreases and resembles more that of intronic absence (C2-DMSO). Indeed, boxes indicate the cells with highest green and red fluorescence, reaching higher percentages with TG003 in a dose dependent manner when the intron is present, and the highest when it is absent. Intron mediated expression enhancement and noise are phenomena that were already described (27) and that can be perfectly exploited in the context of immune gene expression as a quantitative amplifier of rapid transcriptional bursts.

## Discussion

### Changes in CLK1 expression in response to IFN stimulation

When investigating how CLK1 levels are affected in the context of the innate immune response, we identified DAS events that were validated by RT-qPCR and western blot. These events showed an increase in non-coding CLK1 mRNA isoforms after IFN treatment, suggesting a remodeling of the CLK1 splicing pattern (Fig. 2). However, we cannot rule out a possible stimuli-triggered differential stability of the distinct isoforms, a matter that awaits further investigation.

Given that the decrease in the coding isoform is observed in both conditions, viral infection and IFN treatment, it could be hypothesized that this is a response triggered by the host’s immune system and not by the virus itself. However, previous studies from different laboratories, including ours, have shown that the viral protein NS5 interacts with spliceosomal complexes, particularly with components of the U5 snRNP, interfering with host cell splicing and modulating alternative splicing patterns (8). On the other hand, NS5 has also been reported to promote the proteasomal degradation of the splicing factor RBM10, a crucial regulator of the splicing of antiviral pre-mRNAs such as SAT1. Loss of RBM10 leads to changes in SAT1 splicing pattern that reduces the antiviral response, decreasing the production of IFN and proinflammatory cytokines, which favors viral replication (28). Taken together, these findings suggest that both the innate immune response and viral components are involved in the remodeling of cellular splicing in contexts of cellular stress arising from viral infection.

### Silencing and over-expression of CLK1

The CRISPRi system yielded a significant CLK1 silencing, validated by RT-qPCR and western blot. In parallel, the use of CRISPRa induced over-expression of the gene, validated at the RNA and protein levels (Fig 4). However, in the latter case, the cell line showed a severe growth phenotype, hindering its long-term maintenance. In this context, over-expression using a vector was expected to be more efficient. The difficulty in manipulating CLK1 expression may be related to its key role in critical cellular processes, such as cell cycle regulation, among others. CLK1 also auto-regulates its expression through alternative splicing of its own pre-mRNA, by exon 4 skipping and intron retention, generating a feedback loop where high expression of the coding isoform induces the expression of non-coding ones (22). This fine-tuning is fundamental for maintaining adequate CLK1 levels and makes effective silencing or over-expression difficult to achieve. In this regard, previous studies have shown that CLK1 alteration affects cell cycle and reduced proliferation rates, and its over-expression is associated with various types of cancer. These findings reveal an extensive CLK1-dependent alternative splicing program, intimately linked to cell cycle control (29).

### Effect of modulating CLK1 on Immune Response Genes

Silencing CLK1 appears to enhance the action of IFN on the onset of the innate immune response. We observed that the combination of IFN treatment and CLK1 down-regulation using CRISPRi robustly induces the expression of interferon-stimulated genes (ISGs). In the case of ISG15, CLK1 silencing alone is sufficient to significantly increase its expression. These results suggest that CLK1 may be acting as a negative modulator of the IFN pathway, either directly or indirectly, and that its inhibition favors enhanced ISG expression.

CLK1 has been described as a regulator of the alternative splicing of detained introns, a mechanism that can rapidly modulate gene expression in response to cellular stimuli (30). In our case, this could be the addition of IFN or infection with DENV. CLK1 repression could then be favoring the maturation of certain transcripts that, under normal conditions, would remain retained or improperly processed. In this context, it is possible that its inhibition releases restrictions on the splicing of mRNAs involved in the antiviral response. It could be, then, that ISGs maintain their introns retained and that the decrease in CLK1 combined with IFN induction leads to the removal of these introns.

When analyzing RNA-seq data, we were intrigued to evaluate what was happening with ISGs and observed an increase in those IFIT1 and ISG15 isoforms that contain a single intron. This could be consistent with analyses that propose an “economy of expression,” meaning that splicing efficiency increases as transcript levels increase (31). This could indicate the possibility of finer regulation by modulating not only the transcriptional but also the splicing machinery, postulating CLK1 as a node from which it is possible to regulate the function of various splicing factors. Even though our results with the intron-containing reporter suggest that TG003 induces ISG intron mediated expression, further studies are needed to elucidate whether CLK1 acts by modulating the activation, processing and/or stability of transcripts involved in the antiviral response; particularly considering that the GFP transcriptional reporter without any intron (Fig 5) also proved to be affected by TG003 and CLK1 protein levels. IME (Intron Mediated Enhancement) was previously shown to yield fast transcriptional induction, high mRNA output and efficient nuclear export and translation (27), all layers that immune response, especially antiviral, demands. In this case, it is tempting to speculate that CLK1 could exert effects at different layers of ISG expression, including transcription and splicing, which are indeed intrinsically coupled. In this context, it was reported that SRSF7, a member of the SR family of splicing factors and a CLK substrate, facilitates transcription of IRF7 (interferon Regulatory Transcription Factor 7), enhancing innate immunity (32).

### TG003 as a CLK1 inhibitor

TG003 has been characterized as a selective inhibitor of CLK1 and CLK4 (IC50 of 15–20 nM), with a lesser effect on CLK2 (IC50 of 200 nM). This compound has been shown to inhibit SRSF1-dependent splicing of β-globin pre-mRNA *in vitro* by suppressing CLK1-mediated phosphorylation. Furthermore, TG003 blocks the phosphorylation of serine/arginine-rich (SR) proteins, prevents the disassembly of nuclear speckles (structures that act as reservoirs of splicing factors), and disrupts CLK1-dependent alternative splicing in mammalian cells (33).

Figure 5 shows that the inhibitor TG003 does not, by itself, induce the expression of ISGs, unlike what is observed with CLK1 silencing using CRISPRi, where, for example, ISG15 showed increased expression in the absence of IFN stimulation. However, when TG003 is combined with IFN, a significantly greater induction of ISGs such as ISG15 and IFIT1 is observed compared to treatment with IFN alone.This result suggests that both the reduction of CLK1 mRNA levels and the inhibition of its kinase activity affect the expression of ISGs. While this indicates a possible functional connection between CLK1 and the IFN pathway, further investigation would be required to dissect the underlying mechanism.

Previous studies report that SRSF1, a well characterized CLK1 substrate, is extensively involved in the regulation of splicing events directly related to the immune response (34). Another CLK1 substrate, Aurora B, exerts cell cycle-related functions, and its modulation could, in turn, impact the immune response. Nevertheless, it is worth mentioning that the CLK1 phospho-null mutant, that was shown in this study to enhance the induction of the innate immune response, was previously reported to diminish the phosphorylation levels of SRSF1, SRSF2, SRSF4, SRSF5 and SRSF6 (35,36).

On the other hand, a broader and more specific panel of inhibitors could also be tested, given that TG003 has an effect on other CLKs, particularly CLK4 (35). For example, CLK1-IN-1 has been shown to be a molecule designed to more specifically inhibit the CLK1 kinase. With an IC50 of only 2 nM, this means that very little is needed to block kinase activity (37,38). Currently, a significant global effort is invested to develop specific and pharmacologically effective inhibitors of splicing factor kinases, including CLKs, for various clinically viable cancer therapies.

### Effect of the inhibitor TG003 on viral replication

We observed that, when cells were treated with TG003 and infected with DENV-2, viral replication was significantly reduced. This result suggests that pharmacological inhibition of CLKs by TG003 negatively impacts viral replication, leading to the conclusion that CLKs activity is necessary for efficient DENV replication.

The inhibition of splicing factor kinases has already been reported as a promising antiviral strategy. For example, harmine, an inhibitor of SR protein kinases, was found to reduce the levels of various HIV-1 proteins (Env gp160, Gag p55 and p41, and Tat p16 and p14) (39). In the case of influenza A virus, cells treated with the inhibitor TG003 showed a reduction in the replication of the A/WSN/33 strain. Based on this finding, a broader panel of inhibitors was evaluated (21,40). Remarkably, CLK1 was spotted as a chikungunya (CHIKV) proviral hit in an RNAi screen and a few inhibitors as well as CLK1 KO were shown to attenuate CHIKV replication (40). This approach is particularly relevant to be considered for its application in the context of DENV infection.

The obtained results open several avenues for further investigation into the role of CLK1 in regulating the immune response. Exposing cells to individual viral components, such as DENV proteins or synthetic viral RNAs analogs, and other immune stimuli such as bacterial infection, could help to more precisely describe the underlying mechanism. While this manuscript was in preparation, it was reported that CLK1 alternative splicing is modulated in LPS-stimulated PBMCs (41), reinforcing the relevance of CLK1 alternative splicing program in the activation of the innate immunity.

## Acknowledgments

We would like to thank Valeria Buggiano, Mariano Lopez and Amaranta Avendaño for technical help. Evelyn Olszanowski for bioinformatics advice. Diego Alvarez, Melina Magalnik, Laureano Bragado, Nicolás Gaioli y Matías Gambaccini for helpful experimental suggestions and thorough discussion. This work was supported by grants from the Agencia Nacional de Investigaciones Científicas y Tecnológicas of Argentina (PICT 2019-00263 and PICT 2020-00198 to A.S.), the University of Buenos Aires, Argentina (20020220200041BA to A.S.), and the Secretaría de Innovación, Ciencia y Tecnología (former Ministerio de Ciencia, Tecnología e Innovación) of Argentina (Red Federal de Alto Impacto-CONVE-2023-100766162-APN-MCT to A.S.). A.M.L. is recipient of a doctoral fellowship from the CONICET. B.P. and A.S. are career investigators from CONICET.

## Supplementary Information

**Table S1:** Differentially Alternative Spliced (DAS) genes upon DENV infection of A549 cultured cells.

**Table S2:** Differentially Alternative Spliced genes (DAS) upon IFN treatment of PBMCs.

**Table S3:** List of primers for RT-qPCR and cloning

