## Supplementary information for "Targeting CLK1 by CRISPR tools and pharmacological inhibition modulates the innate immune response and dengue virus replication"

| target | contrast | adj.pval | maxdeltaPS | log2FC | up.down |
|---|---|---|---|---|---|
| ENSG00000013441.17 | mock-denv | 0.00264339 | 0.155211598 | 0.038789044 | up-regulated |
| ENSG00000023171.20 | mock-denv | 2.55E-06 | 0.120745713 | 0.178527562 | up-regulated |
| ENSG00000063854.13 | mock-denv | 0.0002665 | -0.19111822 | 0.140574128 | up-regulated |
| ENSG00000071282.12 | mock-denv | 0.00509495 | -0.1099078 | -0.264156294 | down-regulated |
| ENSG00000074527.13 | mock-denv | 0.00281106 | -0.11693433 | -0.783513161 | down-regulated |
| ENSG00000100219.17 | mock-denv | 0.00834772 | -0.15533418 | -1.340981067 | down-regulated |
| ENSG00000100242.16 | mock-denv | 0.00273714 | 0.179100283 | -0.106624822 | down-regulated |
| ENSG00000101160.15 | mock-denv | 0.00023123 | 0.117686467 | 0.292503183 | up-regulated |
| ENSG00000101298.15 | mock-denv | 0.00472879 | 0.31582955 | -0.555122849 | down-regulated |
| ENSG00000107758.16 | mock-denv | 0.00029337 | 0.194858696 | 0.061434542 | up-regulated |
| ENSG00000109320.13 | mock-denv | 0.0078814 | 0.146831256 | -0.404347792 | down-regulated |
| ENSG00000128567.17 | mock-denv | 0.00178224 | -0.11899086 | 0.0328165 | up-regulated |
| ENSG00000130066.17 | mock-denv | 2.04E-12 | -0.13497511 | -0.568756237 | down-regulated |
| ENSG00000134851.13 | mock-denv | 3.01E-05 | 0.20921529 | -0.119938056 | down-regulated |
| ENSG00000135114.13 | mock-denv | 9.17E-07 | 0.785882172 | -5.785720068 | down-regulated |
| ENSG00000137628.18 | mock-denv | 0.00399663 | 0.21815043 | -2.582439813 | down-regulated |
| ENSG00000140105.18 | mock-denv | 1.20E-06 | -0.13411886 | -1.890517967 | down-regulated |
| ENSG00000142089.17 | mock-denv | 0.0056834 | -0.20200661 | -0.877120004 | down-regulated |
| ENSG00000146733.14 | mock-denv | 0.00753525 | 0.151280022 | -0.443934727 | down-regulated |
| ENSG00000147162.14 | mock-denv | 6.99E-08 | 0.110875712 | -0.701529078 | down-regulated |
| ENSG00000152795.18 | mock-denv | 0.00292391 | 0.103269409 | 0.143405826 | up-regulated |
| ENSG00000155660.11 | mock-denv | 0.00066041 | -0.12750695 | -1.719549635 | down-regulated |
| ENSG00000156504.17 | mock-denv | 0.00435009 | -0.1011573 | 0.268931822 | up-regulated |
| ENSG00000159200.18 | mock-denv | 0.00209709 | -0.32632092 | -0.746481241 | down-regulated |
| ENSG00000159403.18 | mock-denv | 0.00485755 | 0.106355808 | -0.907782559 | down-regulated |
| ENSG00000161671.17 | mock-denv | 0.0035646 | -0.10234804 | 0.288826627 | up-regulated |
| ENSG00000164978.18 | mock-denv | 0.00729082 | -0.19051212 | -0.052485711 | down-regulated |
| ENSG00000171302.17 | mock-denv | 8.98E-05 | 0.104108219 | 0.089081648 | up-regulated |
| ENSG00000177410.13 | mock-denv | 0.00079584 | -0.13549169 | -0.905817361 | down-regulated |
| ENSG00000178105.12 | mock-denv | 1.98E-06 | -0.2392158 | -0.774029257 | down-regulated |
| ENSG00000188153.14 | mock-denv | 0.00264339 | -0.15740362 | 0.09290283 | up-regulated |
| ENSG00000198586.14 | mock-denv | 0.00996733 | 0.101540973 | -0.119091583 | down-regulated |
| ENSG00000205746.9 | mock-denv | 0.00906625 | 0.108045934 | -0.78701789 | down-regulated |
| ENSG00000213928.9 | mock-denv | 0.00336402 | -0.2587259 | -1.52628699 | down-regulated |

| target | contrast | adj.pval | maxdeltaPS | log2FC | up.down |
|---|---|---|---|---|---|
| ENSG00000001461.17 | control-ifn | 8.52E-05 | -0.34603279 | 0.551660829 | up-regulated |
| ENSG00000003056.8 | control-ifn | 0.00027689 | -0.22344022 | -0.04321996 | down-regulated |
| ENSG00000003402.21 | control-ifn | 3.69E-09 | 0.18085915 | -0.80396251 | down-regulated |
| ENSG00000003756.17 | control-ifn | 1.55E-06 | -0.12143407 | -0.93241958 | down-regulated |
| ENSG00000004534.15 | control-ifn | 0.00241265 | -0.16001237 | -0.4508551 | down-regulated |
| ENSG00000005483.22 | control-ifn | 7.42E-07 | 0.10677198 | -0.48764778 | down-regulated |
| ENSG00000006125.18 | control-ifn | 0.00082696 | 0.13967758 | -0.57636097 | down-regulated |
| ENSG00000006194.10 | control-ifn | 4.16E-05 | -0.29130771 | 0.485479407 | up-regulated |
| ENSG00000006576.17 | control-ifn | 0.00020689 | -0.66046523 | 0.35657218 | up-regulated |
| ENSG00000006607.14 | control-ifn | 0.00687915 | 0.48145948 | 0.144888891 | up-regulated |
| ENSG00000006652.15 | control-ifn | 4.65E-06 | 0.22766724 | 0.745994231 | up-regulated |
| ENSG00000009954.11 | control-ifn | 3.35E-05 | 0.31437703 | -0.3347179 | down-regulated |
| ENSG00000010244.19 | control-ifn | 4.65E-06 | 0.15000858 | 0.190347786 | up-regulated |
| ENSG00000011114.15 | control-ifn | 0.00036669 | -0.17988385 | -0.22448297 | down-regulated |
| ENSG00000011422.12 | control-ifn | 9.20E-05 | -0.15239039 | 1.743153207 | up-regulated |
| ENSG00000013288.9 | control-ifn | 0.00704344 | -0.12375267 | 0.587654768 | up-regulated |
| ENSG00000013306.16 | control-ifn | 0.00136537 | 0.10760022 | 0.294144917 | up-regulated |
| ENSG00000013374.17 | control-ifn | 3.58E-05 | 0.23707441 | -1.54821667 | down-regulated |
| ENSG00000013441.17 | control-ifn | 0.00115535 | -0.11289768 | 0.429455701 | up-regulated |
| ENSG00000014824.14 | control-ifn | 0.00408674 | 0.15748614 | -0.06640293 | down-regulated |
| ENSG00000015475.19 | control-ifn | 9.14E-06 | 0.22768424 | -0.81708174 | down-regulated |
| ENSG00000015676.18 | control-ifn | 0.00045184 | 0.10558654 | -0.25086544 | down-regulated |
| ENSG00000023041.12 | control-ifn | 0.00664728 | -0.17280817 | -0.38754709 | down-regulated |
| ENSG00000023445.16 | control-ifn | 0.00956846 | -0.13424546 | -1.46193445 | down-regulated |
| ENSG00000023734.11 | control-ifn | 0.00020689 | -0.20144588 | 0.483632749 | up-regulated |
| ENSG00000025293.17 | control-ifn | 0.00082696 | -0.2132805 | 1.452460427 | up-regulated |
| ENSG00000026297.17 | control-ifn | 0.00011434 | 0.10281659 | 1.417385979 | up-regulated |
| ENSG00000028277.22 | control-ifn | 0.00730843 | -0.11437856 | 0.113313393 | up-regulated |
| ENSG00000031698.13 | control-ifn | 0.00362181 | -0.15617964 | -0.30894283 | down-regulated |
| ENSG00000034677.13 | control-ifn | 0.00246299 | -0.14217821 | -0.34843148 | down-regulated |
| ENSG00000036054.13 | control-ifn | 1.83E-05 | -0.4368318 | -1.07498693 | down-regulated |
| ENSG00000039123.16 | control-ifn | 2.98E-05 | 0.33477789 | -0.55433167 | down-regulated |
| ENSG00000039523.20 | control-ifn | 4.04E-05 | -0.46462214 | 0.300445544 | up-regulated |
| ENSG00000042088.14 | control-ifn | 0.0083066 | -0.31904047 | 0.178174753 | up-regulated |
| ENSG00000043143.22 | control-ifn | 2.23E-07 | -0.29870727 | -0.77004315 | down-regulated |
| ENSG00000047644.20 | control-ifn | 0.00182593 | -0.27138473 | 0.069872551 | up-regulated |

| | | | | | |
|---|---|---|---|---|---|
| ENSG00000048162.22 | control-ifn | 0.00202582 | -0.36201862 | -0.33729008 | down-regulated |
| ENSG00000051009.11 | control-ifn | 0.00426739 | 0.37964932 | -0.18352776 | down-regulated |
| ENSG00000052749.14 | control-ifn | 0.00417497 | -0.22087778 | 1.219076803 | up-regulated |
| ENSG00000054267.22 | control-ifn | 4.74E-13 | -0.3910646 | -0.38956518 | down-regulated |
| ENSG00000054523.20 | control-ifn | 0.00010843 | -0.44562991 | -0.32054113 | down-regulated |
| ENSG00000054611.14 | control-ifn | 2.18E-06 | -0.24925404 | 0.418392155 | up-regulated |
| ENSG00000054967.13 | control-ifn | 0.00237474 | -0.14013193 | 0.840834933 | up-regulated |
| ENSG00000055044.11 | control-ifn | 0.00086044 | -0.14041975 | -0.0062602 | down-regulated |
| ENSG00000055483.20 | control-ifn | 0.00402327 | -0.30781015 | -0.06434229 | down-regulated |
| ENSG00000056586.16 | control-ifn | 0.00045184 | 0.50133823 | 0.33302591 | up-regulated |
| ENSG00000057704.13 | control-ifn | 0.00956846 | -0.27667193 | 1.242731229 | up-regulated |
| ENSG00000058272.19 | control-ifn | 0.0018164 | 0.12456954 | -0.70129458 | down-regulated |
| ENSG00000058729.11 | control-ifn | 0.00130878 | 0.25117611 | 0.048492564 | up-regulated |
| ENSG00000059377.18 | control-ifn | 0.00011434 | 0.27493903 | 1.796290171 | up-regulated |
| ENSG00000059588.10 | control-ifn | 0.00038547 | -0.32050287 | -1.279855 | down-regulated |
| ENSG00000060069.18 | control-ifn | 0.0001976 | -0.37885117 | -0.45828348 | down-regulated |
| ENSG00000060971.19 | control-ifn | 0.00161413 | -0.13489114 | 0.587257541 | up-regulated |
| ENSG00000061987.16 | control-ifn | 0.00060072 | -0.12384176 | -0.91952716 | down-regulated |
| ENSG00000062598.18 | control-ifn | 0.0081093 | -0.16393718 | -0.67562688 | down-regulated |
| ENSG00000062716.13 | control-ifn | 0.00049315 | -0.11676074 | 0.455191703 | up-regulated |
| ENSG00000064932.16 | control-ifn | 0.0001976 | -0.23313169 | -1.25798687 | down-regulated |
| ENSG00000065029.15 | control-ifn | 4.62E-05 | 0.12883358 | -0.29749646 | down-regulated |
| ENSG00000065526.12 | control-ifn | 0.00072083 | 0.11531685 | 0.151303387 | up-regulated |
| ENSG00000065675.16 | control-ifn | 0.00849358 | 0.16695743 | 0.257254378 | up-regulated |
| ENSG00000065883.17 | control-ifn | 0.00852625 | 0.29392955 | 0.306492438 | up-regulated |
| ENSG00000066136.21 | control-ifn | 0.00165599 | -0.16187877 | -0.04290664 | down-regulated |
| ENSG00000066379.15 | control-ifn | 9.33E-05 | -0.36858119 | -0.42851611 | down-regulated |
| ENSG00000066422.6 | control-ifn | 0.00064551 | 0.16812475 | 0.202249193 | up-regulated |
| ENSG00000066777.9 | control-ifn | 0.00286033 | -0.11402101 | -0.4440246 | down-regulated |
| ENSG00000067057.18 | control-ifn | 0.00574133 | 0.16409204 | -0.87014601 | down-regulated |
| ENSG00000067064.13 | control-ifn | 0.0001976 | -0.1263088 | 0.991767318 | up-regulated |
| ENSG00000067066.17 | control-ifn | 0.00028913 | 0.1773919 | -1.26335971 | down-regulated |
| ENSG00000067167.8 | control-ifn | 0.00444198 | -0.10122049 | -0.07460865 | down-regulated |
| ENSG00000067248.11 | control-ifn | 0.00256934 | -0.28877161 | 0.065106728 | up-regulated |
| ENSG00000067369.14 | control-ifn | 0.00821016 | -0.25296131 | -1.33137685 | down-regulated |
| ENSG00000067900.8 | control-ifn | 0.00895613 | -0.15194672 | -0.16445604 | down-regulated |
| ENSG00000068028.18 | control-ifn | 0.0065211 | -0.25141785 | 1.102045847 | up-regulated |
| ENSG00000068745.15 | control-ifn | 0.00622179 | 0.2082251 | 0.164701691 | up-regulated |
| ENSG00000068831.19 | control-ifn | 0.00356006 | -0.10725938 | 0.649257442 | up-regulated |

| | | | | | |
|---|---|---|---|---|---|
| ENSG00000069998.12 | control-ifn | 0.00074441 | 0.13319817 | 0.932052924 | up-regulated |
| ENSG00000070423.18 | control-ifn | 0.00620649 | 0.13331732 | 0.865527041 | up-regulated |
| ENSG00000070476.15 | control-ifn | 0.00045626 | 0.25902059 | 0.526391843 | up-regulated |
| ENSG00000070501.12 | control-ifn | 0.00039321 | -0.35691648 | -0.13495854 | down-regulated |
| ENSG00000070614.15 | control-ifn | 0.00079524 | -0.30559585 | 0.430204357 | up-regulated |
| ENSG00000070831.18 | control-ifn | 0.00435856 | -0.18197113 | 0.640015273 | up-regulated |
| ENSG00000071051.14 | control-ifn | 0.00674954 | -0.28506935 | 1.132653697 | up-regulated |
| ENSG00000071073.13 | control-ifn | 6.33E-06 | -0.30906141 | 1.551360657 | up-regulated |
| ENSG00000071127.17 | control-ifn | 6.09E-05 | -0.12267094 | -0.45278233 | down-regulated |
| ENSG00000071243.16 | control-ifn | 2.17E-05 | -0.10737787 | 0.793001546 | up-regulated |
| ENSG00000072756.17 | control-ifn | 0.00202297 | 0.32928486 | -1.08126169 | down-regulated |
| ENSG00000072858.11 | control-ifn | 0.00052664 | -0.31795573 | -0.4162009 | down-regulated |
| ENSG00000073921.18 | control-ifn | 0.00213424 | -0.21430885 | -0.25365479 | down-regulated |
| ENSG00000075413.19 | control-ifn | 0.00302249 | 0.14579498 | -0.11082021 | down-regulated |
| ENSG00000075420.13 | control-ifn | 0.00461808 | 0.25644992 | -2.31376966 | down-regulated |
| ENSG00000077150.20 | control-ifn | 0.00349852 | -0.18251229 | -1.13667336 | down-regulated |
| ENSG00000077235.18 | control-ifn | 1.12E-05 | 0.30315111 | 0.383149276 | up-regulated |
| ENSG00000077420.16 | control-ifn | 3.29E-05 | -0.15582816 | 0.458264697 | up-regulated |
| ENSG00000077721.17 | control-ifn | 0.00165905 | -0.24452167 | -0.58422105 | down-regulated |
| ENSG00000078589.13 | control-ifn | 0.00169635 | -0.16721144 | -0.54046397 | down-regulated |
| ENSG00000078808.19 | control-ifn | 0.00664996 | 0.1071225 | -0.24639028 | down-regulated |
| ENSG00000079277.22 | control-ifn | 0.00058877 | -0.26371463 | -0.64686597 | down-regulated |
| ENSG00000079332.15 | control-ifn | 5.29E-05 | 0.19965341 | -0.15669543 | down-regulated |
| ENSG00000079805.19 | control-ifn | 1.61E-08 | 0.22149216 | 0.073462376 | up-regulated |
| ENSG00000080298.16 | control-ifn | 7.39E-05 | -0.47650203 | 1.217041704 | up-regulated |
| ENSG00000080822.17 | control-ifn | 1.55E-06 | -0.21800681 | -1.22301246 | down-regulated |
| ENSG00000081019.14 | control-ifn | 0.00405011 | 0.26061687 | -0.04898627 | down-regulated |
| ENSG00000081320.11 | control-ifn | 2.26E-06 | -0.33782739 | 1.56656595 | up-regulated |
| ENSG00000083099.11 | control-ifn | 0.00026747 | 0.19479059 | 0.306216234 | up-regulated |
| ENSG00000083123.15 | control-ifn | 0.00077552 | 0.3650506 | 1.199061217 | up-regulated |
| ENSG00000083223.18 | control-ifn | 0.00839558 | 0.24032615 | -0.02795104 | down-regulated |
| ENSG00000083312.19 | control-ifn | 0.00577211 | -0.108236 | -0.95524841 | down-regulated |
| ENSG00000083457.12 | control-ifn | 4.56E-07 | 0.26359838 | 0.515671056 | up-regulated |
| ENSG00000083838.17 | control-ifn | 0.00145416 | -0.54485941 | 0.170566501 | up-regulated |
| ENSG00000083896.13 | control-ifn | 0.00252522 | 0.10396243 | -0.07501443 | down-regulated |
| ENSG00000084234.18 | control-ifn | 0.00574133 | 0.2338316 | 1.596575856 | up-regulated |
| ENSG00000084623.12 | control-ifn | 0.00980265 | -0.15075345 | 0.62029413 | up-regulated |
| ENSG00000085224.23 | control-ifn | 0.00034731 | -0.31136263 | 0.536572051 | up-regulated |
| ENSG00000085415.16 | control-ifn | 0.0015688 | -0.170191 | -0.83078359 | down-regulated |

| | | | | | |
|---|---|---|---|---|---|
| ENSG00000087191.13 | control-ifn | 0.00355283 | 0.11238059 | 0.120026911 | up-regulated |
| ENSG00000087206.17 | control-ifn | 0.00530421 | 0.22755303 | 0.906386813 | up-regulated |
| ENSG00000087589.17 | control-ifn | 0.00023476 | -0.37814195 | 0.038860504 | up-regulated |
| ENSG00000088812.18 | control-ifn | 8.24E-05 | 0.36501489 | 0.187133096 | up-regulated |
| ENSG00000088832.18 | control-ifn | 0.00064548 | -0.11532732 | 1.473226062 | up-regulated |
| ENSG00000089057.15 | control-ifn | 0.00224514 | 0.11541741 | -1.84161724 | down-regulated |
| ENSG00000089094.20 | control-ifn | 0.00926557 | -0.12323106 | -0.46789731 | down-regulated |
| ENSG00000089351.15 | control-ifn | 0.00526582 | 0.12025282 | -0.12249681 | down-regulated |
| ENSG00000089737.18 | control-ifn | 3.33E-07 | 0.14396201 | -0.39971081 | down-regulated |
| ENSG00000089916.18 | control-ifn | 1.30E-05 | 0.14916462 | -0.41520009 | down-regulated |
| ENSG00000090263.16 | control-ifn | 0.00774385 | 0.16944031 | 0.036200674 | up-regulated |
| ENSG00000091527.17 | control-ifn | 7.39E-05 | 0.15473463 | 0.382452782 | up-regulated |
| ENSG00000092330.18 | control-ifn | 5.71E-05 | -0.10262434 | -0.54697127 | down-regulated |
| ENSG00000092931.12 | control-ifn | 0.00218437 | 0.13092681 | 0.013888514 | up-regulated |
| ENSG00000094631.21 | control-ifn | 0.00536239 | -0.13815085 | -0.83426266 | down-regulated |
| ENSG00000094841.14 | control-ifn | 0.00641887 | -0.28035574 | 0.468155268 | up-regulated |
| ENSG00000095303.17 | control-ifn | 0.00446324 | 0.23689295 | 1.087657739 | up-regulated |
| ENSG00000095485.18 | control-ifn | 0.00441622 | 0.14686537 | -0.67367795 | down-regulated |
| ENSG00000095794.20 | control-ifn | 0.00187033 | 0.15241386 | -1.0880486 | down-regulated |
| ENSG00000097007.19 | control-ifn | 0.00441622 | -0.21579462 | 0.462780549 | up-regulated |
| ENSG00000099326.9 | control-ifn | 0.00035033 | 0.26821744 | 0.318593895 | up-regulated |
| ENSG00000099381.19 | control-ifn | 0.00631392 | -0.14983725 | 0.66748139 | up-regulated |
| ENSG00000099968.18 | control-ifn | 6.84E-06 | -0.21297005 | -0.22409684 | down-regulated |
| ENSG00000100038.20 | control-ifn | 0.00010829 | 0.13360256 | -0.36284315 | down-regulated |
| ENSG00000100226.16 | control-ifn | 4.85E-07 | -0.12650233 | -1.53985228 | down-regulated |
| ENSG00000100227.19 | control-ifn | 5.98E-09 | 0.28012452 | -0.39822685 | down-regulated |
| ENSG00000100241.22 | control-ifn | 0.00424524 | -0.10342043 | -0.71629297 | down-regulated |
| ENSG00000100242.16 | control-ifn | 1.04E-11 | 0.1568765 | 0.045209893 | up-regulated |
| ENSG00000100422.14 | control-ifn | 0.00371702 | 0.20940417 | 0.088774256 | up-regulated |
| ENSG00000100461.18 | control-ifn | 5.35E-05 | -0.21351964 | -0.2107213 | down-regulated |
| ENSG00000100519.12 | control-ifn | 0.00010507 | -0.21809574 | -0.53837746 | down-regulated |
| ENSG00000100596.8 | control-ifn | 0.00371702 | -0.35489945 | 1.01909172 | up-regulated |
| ENSG00000100731.16 | control-ifn | 0.0002733 | -0.15649916 | -0.14289184 | down-regulated |
| ENSG00000100796.18 | control-ifn | 0.00186046 | -0.12112323 | -0.06521627 | down-regulated |
| ENSG00000100815.12 | control-ifn | 3.95E-05 | 0.51481055 | -0.37023456 | down-regulated |
| ENSG00000100888.15 | control-ifn | 1.61E-07 | -0.1739668 | -0.36471646 | down-regulated |
| ENSG00000100982.12 | control-ifn | 1.74E-05 | 0.50934868 | 0.883029028 | up-regulated |
| ENSG00000101109.12 | control-ifn | 0.00092588 | 0.12794539 | 0.00914598 | up-regulated |
| ENSG00000101158.15 | control-ifn | 0.00034731 | 0.12440999 | 0.012165579 | up-regulated |

| | | | | | |
|---|---|---|---|---|---|
| ENSG00000101347.11 | control-ifn | 1.18E-12 | -0.18543318 | -0.40864023 | down-regulated |
| ENSG00000101557.15 | control-ifn | 8.89E-06 | -0.18216147 | -0.86582504 | down-regulated |
| ENSG00000101654.18 | control-ifn | 0.00598778 | 0.11212696 | 0.08644463 | up-regulated |
| ENSG00000101935.12 | control-ifn | 0.00706059 | 0.34023257 | 0.658996748 | up-regulated |
| ENSG00000101940.18 | control-ifn | 1.20E-05 | -0.15609124 | 0.809375429 | up-regulated |
| ENSG00000102030.16 | control-ifn | 2.35E-05 | 0.14185716 | 0.43210888 | up-regulated |
| ENSG00000102119.12 | control-ifn | 8.89E-06 | -0.16090314 | 0.055861964 | up-regulated |
| ENSG00000102393.14 | control-ifn | 0.00042955 | -0.36367303 | -0.68516603 | down-regulated |
| ENSG00000102401.20 | control-ifn | 0.00312193 | 0.26442157 | -1.61389728 | down-regulated |
| ENSG00000102409.10 | control-ifn | 0.0006883 | -0.26960015 | 0.541122244 | up-regulated |
| ENSG00000102531.16 | control-ifn | 0.00405916 | -0.29229007 | -0.70001017 | down-regulated |
| ENSG00000102710.21 | control-ifn | 0.00367173 | -0.12447408 | -0.2439868 | down-regulated |
| ENSG00000102804.15 | control-ifn | 0.00687915 | 0.11576145 | -0.27097562 | down-regulated |
| ENSG00000102805.16 | control-ifn | 0.0058082 | -0.20087517 | -0.19305056 | down-regulated |
| ENSG00000103051.20 | control-ifn | 2.09E-05 | 0.32360388 | 0.230904245 | up-regulated |
| ENSG00000103126.15 | control-ifn | 0.00107073 | 0.14435753 | 0.649690607 | up-regulated |
| ENSG00000103194.16 | control-ifn | 9.42E-05 | -0.38152464 | 0.521301068 | up-regulated |
| ENSG00000103197.18 | control-ifn | 0.00126948 | -0.15769856 | -0.67484596 | down-regulated |
| ENSG00000103245.14 | control-ifn | 0.00830908 | -0.31618297 | -0.28238815 | down-regulated |
| ENSG00000103326.12 | control-ifn | 2.04E-05 | 0.12511199 | 0.175806334 | up-regulated |
| ENSG00000103335.22 | control-ifn | 0.00100192 | -0.15641599 | 1.28006463 | up-regulated |
| ENSG00000103429.11 | control-ifn | 0.00131432 | 0.27108932 | 0.796838115 | up-regulated |
| ENSG00000103591.13 | control-ifn | 0.00052317 | -0.15595289 | -0.53031231 | down-regulated |
| ENSG00000104043.15 | control-ifn | 0.00590349 | -0.34296587 | -1.66907525 | down-regulated |
| ENSG00000104164.12 | control-ifn | 0.00046194 | -0.21808813 | 0.855152982 | up-regulated |
| ENSG00000104205.16 | control-ifn | 0.00033964 | -0.32748501 | -0.91299416 | down-regulated |
| ENSG00000104320.14 | control-ifn | 0.00042354 | 0.13573744 | -2.22410899 | down-regulated |
| ENSG00000104447.13 | control-ifn | 0.00609885 | 0.30238136 | 0.420256729 | up-regulated |
| ENSG00000104915.15 | control-ifn | 0.00706059 | 0.19288512 | 2.079284466 | up-regulated |
| ENSG00000104946.14 | control-ifn | 1.11E-08 | 0.26007268 | -1.44889841 | down-regulated |
| ENSG00000105393.16 | control-ifn | 0.00911208 | 0.10254127 | 0.217599999 | up-regulated |
| ENSG00000105643.11 | control-ifn | 0.00032772 | -0.14058474 | -0.42141917 | down-regulated |
| ENSG00000105662.16 | control-ifn | 0.00774385 | -0.45835658 | 0.410062538 | up-regulated |
| ENSG00000105778.20 | control-ifn | 0.00267615 | -0.2236994 | -0.32745046 | down-regulated |
| ENSG00000105879.12 | control-ifn | 5.57E-06 | -0.32428963 | -0.0081645 | down-regulated |
| ENSG00000105953.16 | control-ifn | 0.00026498 | -0.13970688 | -0.0978608 | down-regulated |
| ENSG00000106003.14 | control-ifn | 3.42E-06 | -0.12633422 | -0.81358415 | down-regulated |
| ENSG00000106052.14 | control-ifn | 0.00490079 | -0.14860711 | 0.193616883 | up-regulated |
| ENSG00000106211.10 | control-ifn | 2.21E-09 | -0.61137306 | -0.28377676 | down-regulated |

| | | | | | |
|---|---|---|---|---|---|
| ENSG00000106245.11 | control-ifn | 0.00942716 | 0.18133502 | 0.518218522 | up-regulated |
| ENSG00000106392.11 | control-ifn | 0.00498841 | -0.20291114 | -0.97396661 | down-regulated |
| ENSG00000106443.17 | control-ifn | 0.00011334 | -0.28923418 | 0.061438167 | up-regulated |
| ENSG00000106554.13 | control-ifn | 0.0021403 | -0.1724768 | 1.20793182 | up-regulated |
| ENSG00000106603.20 | control-ifn | 1.53E-05 | -0.11940059 | 0.441723409 | up-regulated |
| ENSG00000106617.15 | control-ifn | 3.48E-07 | -0.21597687 | -0.15308795 | down-regulated |
| ENSG00000106682.16 | control-ifn | 5.87E-10 | -0.34111954 | 0.68316084 | up-regulated |
| ENSG00000106785.15 | control-ifn | 0.00030902 | -0.16783134 | -2.20549936 | down-regulated |
| ENSG00000106829.20 | control-ifn | 0.00081713 | 0.11142668 | -0.74767013 | down-regulated |
| ENSG00000107077.19 | control-ifn | 0.00362945 | -0.20667642 | 0.821817677 | up-regulated |
| ENSG00000107185.10 | control-ifn | 0.00270272 | 0.15772446 | -0.38379947 | down-regulated |
| ENSG00000107262.26 | control-ifn | 0.00020758 | 0.17248104 | -0.88363174 | down-regulated |
| ENSG00000107372.13 | control-ifn | 9.09E-07 | -0.1691149 | 0.863531024 | up-regulated |
| ENSG00000107651.14 | control-ifn | 0.00038935 | 0.48618662 | -0.32802496 | down-regulated |
| ENSG00000107938.18 | control-ifn | 1.05E-08 | -0.31430951 | 0.171232727 | up-regulated |
| ENSG00000107951.15 | control-ifn | 0.00581098 | 0.34292903 | 1.245226537 | up-regulated |
| ENSG00000107960.11 | control-ifn | 0.00094811 | 0.17017768 | 1.289364287 | up-regulated |
| ENSG00000107968.10 | control-ifn | 0.00046245 | -0.22891189 | 1.09167142 | up-regulated |
| ENSG00000108219.15 | control-ifn | 0.00611841 | 0.15522372 | 0.574587146 | up-regulated |
| ENSG00000108312.15 | control-ifn | 0.00092467 | 0.12048965 | -0.34231552 | down-regulated |
| ENSG00000108510.10 | control-ifn | 0.00464059 | 0.21656985 | -0.75118023 | down-regulated |
| ENSG00000108671.11 | control-ifn | 0.00072675 | 0.12435135 | -1.19000443 | down-regulated |
| ENSG00000108946.16 | control-ifn | 4.94E-05 | 0.17378492 | -0.17776989 | down-regulated |
| ENSG00000109046.15 | control-ifn | 0.00595544 | -0.11811469 | -0.56471112 | down-regulated |
| ENSG00000109606.13 | control-ifn | 0.0001984 | -0.21748695 | -0.13243211 | down-regulated |
| ENSG00000109670.16 | control-ifn | 0.00028827 | 0.29915859 | 0.478409225 | up-regulated |
| ENSG00000109689.19 | control-ifn | 0.00032098 | 0.13581925 | 0.151271674 | up-regulated |
| ENSG00000109787.13 | control-ifn | 6.39E-07 | 0.58071767 | 1.330953498 | up-regulated |
| ENSG00000109790.18 | control-ifn | 0.00556865 | -0.11287848 | -0.8520725 | down-regulated |
| ENSG00000109929.10 | control-ifn | 0.00287918 | -0.24208103 | -0.21572017 | down-regulated |
| ENSG00000110046.13 | control-ifn | 0.00506415 | -0.19419296 | 0.339231539 | up-regulated |
| ENSG00000110066.15 | control-ifn | 9.20E-05 | -0.1806659 | -0.32762933 | down-regulated |
| ENSG00000110172.12 | control-ifn | 3.00E-06 | -0.2198963 | -1.77189413 | down-regulated |
| ENSG00000110315.7 | control-ifn | 0.0035935 | 0.12963578 | 2.061863552 | up-regulated |
| ENSG00000110324.12 | control-ifn | 0.00315408 | 0.11435911 | 0.065046846 | up-regulated |
| ENSG00000110367.14 | control-ifn | 0.00140223 | -0.1199759 | -0.18549107 | down-regulated |
| ENSG00000110429.14 | control-ifn | 0.00117619 | -0.23454163 | -0.29558616 | down-regulated |
| ENSG00000110848.8 | control-ifn | 0.00522103 | 0.12487112 | 3.261377795 | up-regulated |
| ENSG00000110917.8 | control-ifn | 0.00408674 | -0.28211985 | -0.70495453 | down-regulated |

| | | | | | |
|---|---|---|---|---|---|
| ENSG00000110931.19 | control-ifn | 0.00378379 | 0.19366819 | -0.53281194 | down-regulated |
| ENSG00000110934.13 | control-ifn | 0.00056821 | 0.1181734 | 1.276881607 | up-regulated |
| ENSG00000111300.10 | control-ifn | 0.00048412 | -0.27950848 | -1.08988556 | down-regulated |
| ENSG00000111331.14 | control-ifn | 6.15E-05 | 0.11216937 | -4.67425297 | down-regulated |
| ENSG00000111802.14 | control-ifn | 3.07E-05 | -0.25837846 | -0.69040948 | down-regulated |
| ENSG00000111859.17 | control-ifn | 0.00069225 | -0.20764894 | 1.664363048 | up-regulated |
| ENSG00000111913.20 | control-ifn | 0.00082324 | -0.34752324 | 0.415303807 | up-regulated |
| ENSG00000112200.17 | control-ifn | 0.00014086 | 0.20013084 | -0.52651182 | down-regulated |
| ENSG00000112592.15 | control-ifn | 0.00785269 | 0.23211261 | -0.56110026 | down-regulated |
| ENSG00000112763.17 | control-ifn | 0.00581852 | -0.18553591 | -0.76834082 | down-regulated |
| ENSG00000112996.11 | control-ifn | 0.00273212 | -0.12550631 | 0.350074074 | up-regulated |
| ENSG00000113163.19 | control-ifn | 0.00380856 | -0.13047713 | 0.342184938 | up-regulated |
| ENSG00000113441.16 | control-ifn | 0.00349947 | -0.16209065 | -1.27105742 | down-regulated |
| ENSG00000113580.15 | control-ifn | 0.0027198 | -0.14397512 | -0.58250216 | down-regulated |
| ENSG00000113648.17 | control-ifn | 0.00195318 | 0.12886611 | 0.657820175 | up-regulated |
| ENSG00000113712.19 | control-ifn | 0.00014066 | 0.22834525 | -0.29799945 | down-regulated |
| ENSG00000113845.10 | control-ifn | 0.00578183 | -0.27075956 | 0.042584618 | up-regulated |
| ENSG00000114098.18 | control-ifn | 0.00038315 | -0.1464347 | 0.149358934 | up-regulated |
| ENSG00000114127.11 | control-ifn | 0.00059898 | 0.1278425 | -2.28222882 | down-regulated |
| ENSG00000114416.18 | control-ifn | 8.31E-05 | 0.13445628 | 0.407064437 | up-regulated |
| ENSG00000114423.23 | control-ifn | 0.00288484 | -0.26780044 | -0.27513387 | down-regulated |
| ENSG00000114439.19 | control-ifn | 0.00203814 | -0.26286516 | 0.421005106 | up-regulated |
| ENSG00000114503.11 | control-ifn | 0.00397777 | 0.10316728 | -0.56625738 | down-regulated |
| ENSG00000114796.16 | control-ifn | 0.00081047 | 0.19847591 | 0.791339876 | up-regulated |
| ENSG00000114867.22 | control-ifn | 0.00144601 | -0.19349118 | -0.53427184 | down-regulated |
| ENSG00000114982.19 | control-ifn | 0.00254536 | -0.13313778 | -0.64541243 | down-regulated |
| ENSG00000115267.9 | control-ifn | 6.06E-06 | 0.48009188 | -3.57513665 | down-regulated |
| ENSG00000115325.14 | control-ifn | 0.00198089 | 0.46990566 | -0.47652266 | down-regulated |
| ENSG00000115484.15 | control-ifn | 0.00413896 | -0.22461527 | 0.104035908 | up-regulated |
| ENSG00000115904.15 | control-ifn | 9.23E-09 | 0.35306133 | -0.64951502 | down-regulated |
| ENSG00000115935.18 | control-ifn | 8.08E-13 | 0.21292062 | -0.65581642 | down-regulated |
| ENSG00000116337.20 | control-ifn | 0.00022401 | -0.23473457 | 2.173368828 | up-regulated |
| ENSG00000116478.12 | control-ifn | 0.00201256 | 0.14751802 | 0.220699054 | up-regulated |
| ENSG00000116580.20 | control-ifn | 0.00412828 | 0.17841594 | -0.3757407 | down-regulated |
| ENSG00000116584.22 | control-ifn | 0.00042851 | -0.22302342 | -0.62217814 | down-regulated |
| ENSG00000116679.16 | control-ifn | 2.93E-10 | -0.2098117 | 0.233167892 | up-regulated |
| ENSG00000116685.16 | control-ifn | 0.00246299 | 0.20969195 | 0.391610123 | up-regulated |
| ENSG00000116688.18 | control-ifn | 0.00028031 | -0.2820227 | 0.04845456 | up-regulated |
| ENSG00000116747.13 | control-ifn | 0.00969765 | -0.29573646 | 0.288426336 | up-regulated |

| | | | | | |
|---|---|---|---|---|---|
| ENSG00000116809.12 | control-ifn | 0.00049738 | -0.13295381 | 0.065410783 | up-regulated |
| ENSG00000116830.12 | control-ifn | 3.49E-06 | 0.25213542 | -0.45797372 | down-regulated |
| ENSG00000116871.16 | control-ifn | 0.0005478 | -0.20184048 | -0.31684308 | down-regulated |
| ENSG00000116977.19 | control-ifn | 0.00182593 | 0.11027604 | -0.29298881 | down-regulated |
| ENSG00000116984.15 | control-ifn | 9.53E-05 | 0.19044487 | -1.16668141 | down-regulated |
| ENSG00000117281.16 | control-ifn | 0.00479302 | -0.23694243 | 3.369050972 | up-regulated |
| ENSG00000117298.16 | control-ifn | 0.00020758 | -0.37637363 | -0.4111324 | down-regulated |
| ENSG00000117475.14 | control-ifn | 0.00441622 | 0.24370933 | -2.1502683 | down-regulated |
| ENSG00000117528.14 | control-ifn | 0.00053387 | 0.43460128 | 0.082548095 | up-regulated |
| ENSG00000118058.24 | control-ifn | 0.00025564 | -0.19839136 | -0.48018708 | down-regulated |
| ENSG00000118217.7 | control-ifn | 0.00121577 | 0.15954925 | -0.15348072 | down-regulated |
| ENSG00000118260.15 | control-ifn | 5.74E-06 | -0.14979961 | 0.462960235 | up-regulated |
| ENSG00000118482.12 | control-ifn | 6.58E-05 | -0.121308 | 0.530966167 | up-regulated |
| ENSG00000118503.16 | control-ifn | 3.17E-05 | -0.13061685 | 1.517250471 | up-regulated |
| ENSG00000118507.18 | control-ifn | 8.54E-06 | -0.28728026 | 0.749682205 | up-regulated |
| ENSG00000119231.11 | control-ifn | 0.00030945 | 0.25279625 | -0.70797223 | down-regulated |
| ENSG00000119402.18 | control-ifn | 0.00036221 | -0.13748847 | -0.33485531 | down-regulated |
| ENSG00000119408.17 | control-ifn | 0.00144601 | 0.23761298 | 0.207504351 | up-regulated |
| ENSG00000119866.22 | control-ifn | 0.00047373 | 0.37532124 | 1.043293641 | up-regulated |
| ENSG00000119878.6 | control-ifn | 0.00667188 | -0.24819117 | -0.53090381 | down-regulated |
| ENSG00000120616.16 | control-ifn | 9.23E-09 | -0.24456066 | 0.974964336 | up-regulated |
| ENSG00000120694.20 | control-ifn | 0.00128052 | -0.14288904 | -1.86007343 | down-regulated |
| ENSG00000121067.19 | control-ifn | 0.00385378 | 0.187826 | -0.39283488 | down-regulated |
| ENSG00000121350.16 | control-ifn | 0.00201242 | 0.41497398 | 0.711737221 | up-regulated |
| ENSG00000121741.17 | control-ifn | 5.74E-06 | -0.16653853 | -0.07155267 | down-regulated |
| ENSG00000121895.8 | control-ifn | 0.00029275 | -0.47001485 | 1.042967645 | up-regulated |
| ENSG00000122257.20 | control-ifn | 2.87E-05 | -0.3069701 | 1.64617468 | up-regulated |
| ENSG00000122643.23 | control-ifn | 8.08E-13 | 0.37699881 | -3.56958734 | down-regulated |
| ENSG00000122707.12 | control-ifn | 0.00034731 | -0.39910647 | 0.790634141 | up-regulated |
| ENSG00000122729.19 | control-ifn | 0.00277139 | 0.26377485 | -0.38088174 | down-regulated |
| ENSG00000122783.17 | control-ifn | 3.28E-06 | 0.21389085 | -2.49461978 | down-regulated |
| ENSG00000123136.15 | control-ifn | 4.82E-06 | -0.20646602 | 0.488234108 | up-regulated |
| ENSG00000123200.17 | control-ifn | 0.00011434 | 0.34469522 | -0.32858322 | down-regulated |
| ENSG00000123329.20 | control-ifn | 9.41E-10 | 0.10730422 | 0.955812781 | up-regulated |
| ENSG00000123358.20 | control-ifn | 1.12E-06 | 0.16565127 | 4.436174827 | up-regulated |
| ENSG00000123545.6 | control-ifn | 0.00618868 | 0.19398526 | 0.296968203 | up-regulated |
| ENSG00000123562.18 | control-ifn | 0.00288484 | 0.20127062 | -0.27231364 | down-regulated |
| ENSG00000123983.15 | control-ifn | 0.0013146 | -0.19734928 | -0.01615638 | down-regulated |
| ENSG00000124151.19 | control-ifn | 2.40E-05 | 0.22508917 | -0.52527118 | down-regulated |

| | | | | | |
|---|---|---|---|---|---|
| ENSG00000124406.16 | control-ifn | 0.0003587 | -0.17112759 | -0.76916327 | down-regulated |
| ENSG00000124596.17 | control-ifn | 0.002352 | 0.19389329 | 0.098059451 | up-regulated |
| ENSG00000124731.13 | control-ifn | 0.00122024 | -0.10959062 | 1.806766795 | up-regulated |
| ENSG00000124782.21 | control-ifn | 2.26E-06 | -0.25874336 | 0.569631957 | up-regulated |
| ENSG00000125089.18 | control-ifn | 0.00023476 | -0.28402951 | 0.525881374 | up-regulated |
| ENSG00000125107.19 | control-ifn | 1.25E-07 | 0.21838663 | -0.31073012 | down-regulated |
| ENSG00000125430.9 | control-ifn | 0.00526168 | -0.34095745 | -1.07437455 | down-regulated |
| ENSG00000125454.12 | control-ifn | 0.00340456 | 0.36935086 | -2.57781745 | down-regulated |
| ENSG00000125676.20 | control-ifn | 0.00131992 | -0.1255756 | -0.52806278 | down-regulated |
| ENSG00000125730.18 | control-ifn | 0.00533101 | 0.57736248 | -3.48384116 | down-regulated |
| ENSG00000125741.6 | control-ifn | 0.00632819 | 0.27355991 | -0.07950303 | down-regulated |
| ENSG00000125746.18 | control-ifn | 0.00073095 | 0.2414478 | 0.277742193 | up-regulated |
| ENSG00000125834.13 | control-ifn | 0.00205129 | 0.24764092 | -0.1804335 | down-regulated |
| ENSG00000125871.14 | control-ifn | 0.00792486 | -0.36815836 | 0.201065934 | up-regulated |
| ENSG00000126016.17 | control-ifn | 0.00356006 | 0.3886332 | 0.273857696 | up-regulated |
| ENSG00000126106.14 | control-ifn | 0.00288484 | 0.67641504 | 1.251053329 | up-regulated |
| ENSG00000126107.15 | control-ifn | 0.00405083 | -0.1846809 | -0.88642949 | down-regulated |
| ENSG00000126214.22 | control-ifn | 8.33E-05 | 0.20191513 | -0.64457832 | down-regulated |
| ENSG00000126216.15 | control-ifn | 0.00133868 | -0.18574212 | 0.376976648 | up-regulated |
| ENSG00000126934.15 | control-ifn | 0.00013963 | 0.12532132 | 0.592094206 | up-regulated |
| ENSG00000127452.9 | control-ifn | 0.00048877 | -0.33515296 | 0.138248805 | up-regulated |
| ENSG00000127481.15 | control-ifn | 0.00346255 | 0.2113899 | -0.7524384 | down-regulated |
| ENSG00000127838.15 | control-ifn | 0.00177557 | 0.38260573 | 0.18454432 | up-regulated |
| ENSG00000128271.22 | control-ifn | 3.20E-05 | -0.44382451 | -1.3502569 | down-regulated |
| ENSG00000128284.20 | control-ifn | 6.78E-05 | -0.12847216 | -2.39989916 | down-regulated |
| ENSG00000128294.16 | control-ifn | 0.00281339 | 0.14978827 | 0.155283163 | up-regulated |
| ENSG00000128309.17 | control-ifn | 0.00043705 | 0.16395931 | 2.009347642 | up-regulated |
| ENSG00000128595.17 | control-ifn | 0.00175931 | -0.18079565 | 0.278741653 | up-regulated |
| ENSG00000129071.10 | control-ifn | 0.00649444 | 0.15159956 | 0.105782641 | up-regulated |
| ENSG00000129292.21 | control-ifn | 0.00441622 | -0.12463738 | -0.13535788 | down-regulated |
| ENSG00000129315.11 | control-ifn | 0.00851596 | 0.14209268 | 0.147726279 | up-regulated |
| ENSG00000130244.13 | control-ifn | 0.00213169 | -0.22182784 | 0.481694913 | up-regulated |
| ENSG00000130304.17 | control-ifn | 0.00049315 | 0.27630528 | 2.120086266 | up-regulated |
| ENSG00000130309.12 | control-ifn | 0.0021403 | 0.26914408 | 0.983351188 | up-regulated |
| ENSG00000130313.7 | control-ifn | 0.00200732 | 0.12008672 | 1.381775393 | up-regulated |
| ENSG00000130363.13 | control-ifn | 0.0091275 | -0.25382918 | 0.314685034 | up-regulated |
| ENSG00000130803.15 | control-ifn | 1.32E-07 | -0.46234158 | 0.844395645 | up-regulated |
| ENSG00000130813.18 | control-ifn | 0.00622179 | 0.13874044 | -2.64702959 | down-regulated |
| ENSG00000130844.19 | control-ifn | 1.08E-05 | -0.35279254 | 2.114974021 | up-regulated |

| | | | | | |
|---|---|---|---|---|---|
| ENSG00000130921.9 | control-ifn | 0.00062923 | -0.31808172 | -0.66883222 | down-regulated |
| ENSG00000131018.25 | control-ifn | 0.00017571 | 0.13189027 | -0.47410953 | down-regulated |
| ENSG00000131023.13 | control-ifn | 2.67E-07 | 0.48239467 | 0.44215881 | up-regulated |
| ENSG00000131263.13 | control-ifn | 0.00288508 | 0.21497023 | 0.695037854 | up-regulated |
| ENSG00000131375.10 | control-ifn | 0.00430046 | -0.13098126 | 0.312580885 | up-regulated |
| ENSG00000131408.15 | control-ifn | 1.09E-05 | -0.13681958 | 0.566410329 | up-regulated |
| ENSG00000131446.17 | control-ifn | 0.00569341 | -0.17443903 | -0.02052139 | down-regulated |
| ENSG00000131979.20 | control-ifn | 4.43E-06 | 0.47445216 | -1.36329022 | down-regulated |
| ENSG00000132294.15 | control-ifn | 0.00277899 | 0.15072423 | -1.68296614 | down-regulated |
| ENSG00000132334.17 | control-ifn | 6.22E-07 | -0.22418549 | -0.55102354 | down-regulated |
| ENSG00000132388.13 | control-ifn | 0.00388755 | -0.19539533 | 0.459924985 | up-regulated |
| ENSG00000132463.15 | control-ifn | 0.00802034 | 0.11305996 | 0.266324594 | up-regulated |
| ENSG00000132530.17 | control-ifn | 8.49E-06 | 0.19621418 | -4.5126673 | down-regulated |
| ENSG00000132680.11 | control-ifn | 0.00010317 | -0.32509775 | -1.44500479 | down-regulated |
| ENSG00000132773.12 | control-ifn | 0.00768199 | -0.14683421 | 1.065928457 | up-regulated |
| ENSG00000132950.19 | control-ifn | 0.00028333 | 0.37077684 | 0.056472514 | up-regulated |
| ENSG00000133026.14 | control-ifn | 0.00687915 | 0.4014442 | 2.06264586 | up-regulated |
| ENSG00000133243.10 | control-ifn | 6.52E-08 | 0.39649717 | 0.373058307 | up-regulated |
| ENSG00000133316.16 | control-ifn | 0.00053382 | -0.20450282 | 0.277416298 | up-regulated |
| ENSG00000133619.18 | control-ifn | 0.00938638 | 0.29327803 | 0.104340675 | up-regulated |
| ENSG00000133657.17 | control-ifn | 3.19E-05 | -0.12004334 | -1.92455372 | down-regulated |
| ENSG00000133805.16 | control-ifn | 1.79E-05 | -0.23475922 | -1.78205153 | down-regulated |
| ENSG00000133872.14 | control-ifn | 4.82E-05 | -0.12430132 | 0.126147266 | up-regulated |
| ENSG00000133961.21 | control-ifn | 0.00079444 | 0.19224496 | -0.30746895 | down-regulated |
| ENSG00000133997.12 | control-ifn | 4.71E-05 | -0.19098746 | -0.09266967 | down-regulated |
| ENSG00000134077.16 | control-ifn | 1.09E-05 | 0.35289743 | -0.36477778 | down-regulated |
| ENSG00000134186.12 | control-ifn | 0.00155314 | 0.11331819 | -0.19725421 | down-regulated |
| ENSG00000134242.16 | control-ifn | 0.00424098 | -0.26243008 | -0.32783338 | down-regulated |
| ENSG00000134248.14 | control-ifn | 0.0011141 | -0.18611617 | 0.54762089 | up-regulated |
| ENSG00000134250.21 | control-ifn | 6.11E-05 | -0.252373 | -0.1627531 | down-regulated |
| ENSG00000134294.14 | control-ifn | 4.82E-06 | -0.15045186 | -0.15383141 | down-regulated |
| ENSG00000134321.13 | control-ifn | 6.01E-06 | -0.53284598 | -6.55411193 | down-regulated |
| ENSG00000134686.20 | control-ifn | 0.00080727 | -0.1215861 | 0.898911134 | up-regulated |
| ENSG00000134884.15 | control-ifn | 0.00175511 | -0.21685057 | 0.686044424 | up-regulated |
| ENSG00000135077.10 | control-ifn | 0.00674506 | -0.23112624 | -1.7146466 | down-regulated |
| ENSG00000135090.14 | control-ifn | 0.00441622 | -0.14607934 | -0.25524646 | down-regulated |
| ENSG00000135114.13 | control-ifn | 0.00031007 | -0.14065356 | -4.58239411 | down-regulated |
| ENSG00000135144.8 | control-ifn | 0.00029275 | 0.50841504 | 0.567804055 | up-regulated |
| ENSG00000135250.17 | control-ifn | 0.00475594 | 0.21032645 | 0.83403124 | up-regulated |

| | | | | | |
|---|---|---|---|---|---|
| ENSG00000135269.18 | control-ifn | 0.00098573 | -0.16514901 | 0.314185779 | up-regulated |
| ENSG00000135316.19 | control-ifn | 0.00337424 | -0.14278742 | -0.50816752 | down-regulated |
| ENSG00000135446.17 | control-ifn | 0.00577211 | 0.2433307 | -0.36626873 | down-regulated |
| ENSG00000135637.14 | control-ifn | 0.00890977 | 0.22897519 | -0.53645497 | down-regulated |
| ENSG00000135686.13 | control-ifn | 7.39E-05 | 0.30008877 | -0.1358586 | down-regulated |
| ENSG00000136068.16 | control-ifn | 0.00369229 | -0.23564406 | -0.24258643 | down-regulated |
| ENSG00000136152.15 | control-ifn | 2.33E-05 | -0.39516201 | 1.533804659 | up-regulated |
| ENSG00000136250.12 | control-ifn | 0.00814048 | 0.12187316 | 0.869221213 | up-regulated |
| ENSG00000136273.13 | control-ifn | 0.00050453 | 0.3792089 | 0.403070204 | up-regulated |
| ENSG00000136280.17 | control-ifn | 0.00038074 | -0.34346613 | -0.23873606 | down-regulated |
| ENSG00000136305.12 | control-ifn | 0.00644439 | -0.28793069 | 3.150693226 | up-regulated |
| ENSG00000136381.13 | control-ifn | 0.0011141 | -0.26705329 | -0.26118778 | down-regulated |
| ENSG00000136448.13 | control-ifn | 1.12E-07 | 0.30163915 | 0.042468186 | up-regulated |
| ENSG00000136485.16 | control-ifn | 0.00451093 | -0.42468697 | -0.83312481 | down-regulated |
| ENSG00000136527.19 | control-ifn | 9.09E-07 | 0.16939617 | 0.26458015 | up-regulated |
| ENSG00000136536.15 | control-ifn | 0.00037318 | 0.11230799 | -0.4616072 | down-regulated |
| ENSG00000136689.20 | control-ifn | 0.00553958 | -0.11947503 | -2.98740898 | down-regulated |
| ENSG00000136758.20 | control-ifn | 0.00224665 | -0.13270733 | -0.61824518 | down-regulated |
| ENSG00000136861.19 | control-ifn | 0.00382445 | -0.10662665 | -0.53321264 | down-regulated |
| ENSG00000136878.14 | control-ifn | 0.00271958 | -0.12023331 | -0.21877621 | down-regulated |
| ENSG00000136908.18 | control-ifn | 0.00087371 | -0.21044227 | -0.17786357 | down-regulated |
| ENSG00000136929.13 | control-ifn | 0.00064738 | -1 | -0.0701067 | down-regulated |
| ENSG00000136932.15 | control-ifn | 0.00415131 | 0.25995955 | 0.895428565 | up-regulated |
| ENSG00000136997.21 | control-ifn | 0.00384507 | 0.12622814 | -1.64225368 | down-regulated |
| ENSG00000137075.18 | control-ifn | 0.00110937 | -0.1870887 | 0.22628447 | up-regulated |
| ENSG00000137275.16 | control-ifn | 6.18E-05 | -0.28155841 | -1.34416508 | down-regulated |
| ENSG00000137500.10 | control-ifn | 0.00631392 | -0.15542385 | -0.72287713 | down-regulated |
| ENSG00000137501.18 | control-ifn | 0.00589194 | -0.21717183 | -1.57618189 | down-regulated |
| ENSG00000137770.14 | control-ifn | 0.00019293 | -0.11170076 | 0.05742456 | up-regulated |
| ENSG00000137841.12 | control-ifn | 1.16E-08 | 0.11251946 | 2.570668363 | up-regulated |
| ENSG00000137871.21 | control-ifn | 0.00528123 | -0.10784621 | 0.766165597 | up-regulated |
| ENSG00000138035.15 | control-ifn | 0.00221719 | -0.47598546 | -3.7246822 | down-regulated |
| ENSG00000138442.11 | control-ifn | 0.0017936 | -0.15607504 | -0.30032865 | down-regulated |
| ENSG00000138614.16 | control-ifn | 0.00349947 | 0.4113755 | -0.59340991 | down-regulated |
| ENSG00000138835.23 | control-ifn | 0.00331446 | 0.21417134 | -0.41890451 | down-regulated |
| ENSG00000139163.16 | control-ifn | 0.0004373 | -0.36204825 | -1.16086946 | down-regulated |
| ENSG00000139192.12 | control-ifn | 0.00106467 | 0.29611217 | -0.11008395 | down-regulated |
| ENSG00000139218.18 | control-ifn | 3.50E-07 | -0.23717797 | -0.59688762 | down-regulated |
| ENSG00000139436.22 | control-ifn | 1.39E-06 | 0.1627248 | 0.292794705 | up-regulated |

| | | | | | |
|---|---|---|---|---|---|
| ENSG00000139496.17 | control-ifn | 5.88E-07 | -0.36121385 | -0.7315793 | down-regulated |
| ENSG00000139597.18 | control-ifn | 4.71E-06 | 0.30664899 | 0.52059894 | up-regulated |
| ENSG00000139636.16 | control-ifn | 0.00045438 | -0.10130782 | -0.14554938 | down-regulated |
| ENSG00000139793.20 | control-ifn | 0.00046732 | -0.29152754 | 1.490112741 | up-regulated |
| ENSG00000139842.15 | control-ifn | 0.00289578 | 0.21889918 | -1.2581641 | down-regulated |
| ENSG00000140262.18 | control-ifn | 0.00011434 | 0.29348651 | -0.89386937 | down-regulated |
| ENSG00000140455.17 | control-ifn | 0.00123313 | 0.17875061 | 0.620560528 | up-regulated |
| ENSG00000140987.21 | control-ifn | 0.00808589 | 0.24136724 | -1.13304891 | down-regulated |
| ENSG00000140995.17 | control-ifn | 0.00012252 | 0.17616658 | 1.240643101 | up-regulated |
| ENSG00000141012.13 | control-ifn | 1.28E-05 | 0.13662245 | 0.662144216 | up-regulated |
| ENSG00000141034.10 | control-ifn | 0.00833041 | -0.19003831 | -0.4493007 | down-regulated |
| ENSG00000141298.19 | control-ifn | 0.0018135 | -0.18521415 | 0.87559338 | up-regulated |
| ENSG00000141337.13 | control-ifn | 0.00280111 | 0.46917157 | 0.974686502 | up-regulated |
| ENSG00000141401.12 | control-ifn | 0.00059172 | -0.23157961 | 3.615560461 | up-regulated |
| ENSG00000141447.19 | control-ifn | 0.00062923 | 0.50829714 | 1.149514493 | up-regulated |
| ENSG00000141504.12 | control-ifn | 0.00035558 | -0.152882 | 0.778803873 | up-regulated |
| ENSG00000141510.18 | control-ifn | 0.00182593 | -0.13745778 | -0.61171846 | down-regulated |
| ENSG00000141526.18 | control-ifn | 0.00010829 | -0.14117617 | -2.16768689 | down-regulated |
| ENSG00000141556.22 | control-ifn | 0.00010909 | 0.10648701 | -0.56937101 | down-regulated |
| ENSG00000141562.19 | control-ifn | 0.00514814 | 0.14190504 | -0.36136597 | down-regulated |
| ENSG00000141664.10 | control-ifn | 0.00553958 | 0.12818266 | -2.27088828 | down-regulated |
| ENSG00000141837.22 | control-ifn | 0.00203814 | 0.39795932 | -1.39148292 | down-regulated |
| ENSG00000141867.19 | control-ifn | 0.00890977 | -0.21906885 | -0.2441392 | down-regulated |
| ENSG00000142166.13 | control-ifn | 6.76E-08 | 0.32672021 | -0.33872213 | down-regulated |
| ENSG00000142687.18 | control-ifn | 0.00956846 | 0.10060926 | -0.47059324 | down-regulated |
| ENSG00000142765.18 | control-ifn | 6.15E-05 | 0.12980564 | 1.558144126 | up-regulated |
| ENSG00000142864.15 | control-ifn | 0.0029153 | -0.12010458 | 0.057185828 | up-regulated |
| ENSG00000142867.14 | control-ifn | 0.00424524 | 0.1793659 | 0.202068717 | up-regulated |
| ENSG00000143164.16 | control-ifn | 0.00066657 | 0.26669889 | 0.044995042 | up-regulated |
| ENSG00000143258.17 | control-ifn | 0.00528726 | -0.20508049 | -0.26993182 | down-regulated |
| ENSG00000143344.16 | control-ifn | 0.00034731 | 0.27895804 | -6.07403519 | down-regulated |
| ENSG00000143376.14 | control-ifn | 8.53E-05 | -0.13759507 | 0.291835666 | up-regulated |
| ENSG00000143398.20 | control-ifn | 2.10E-08 | 0.42737846 | -0.5211836 | down-regulated |
| ENSG00000143537.14 | control-ifn | 0.00171995 | 0.2251249 | 1.749758562 | up-regulated |
| ENSG00000143569.19 | control-ifn | 0.00024153 | 0.11132433 | -0.3553547 | down-regulated |
| ENSG00000143669.15 | control-ifn | 1.36E-09 | 0.35028506 | 1.119448473 | up-regulated |
| ENSG00000143756.12 | control-ifn | 0.00764629 | 0.50983947 | 0.511832091 | up-regulated |
| ENSG00000143771.12 | control-ifn | 8.89E-05 | 0.15024439 | 0.34114596 | up-regulated |
| ENSG00000144136.11 | control-ifn | 0.00581031 | -0.11797411 | 0.167355387 | up-regulated |

| | | | | | |
|---|---|---|---|---|---|
| ENSG00000144566.11 | control-ifn | 2.17E-05 | 0.32282062 | -0.5036842 | down-regulated |
| ENSG00000144802.11 | control-ifn | 7.40E-08 | 0.19850611 | 1.506403955 | up-regulated |
| ENSG00000145241.11 | control-ifn | 7.39E-05 | 0.24018181 | 0.27096913 | up-regulated |
| ENSG00000145391.14 | control-ifn | 0.00061416 | -0.38641427 | 2.034857737 | up-regulated |
| ENSG00000145414.9 | control-ifn | 0.00403703 | 0.30648265 | 0.587085296 | up-regulated |
| ENSG00000145740.20 | control-ifn | 0.00013496 | -0.22784958 | -0.48901017 | down-regulated |
| ENSG00000145779.8 | control-ifn | 3.33E-07 | 0.16521899 | -1.10130764 | down-regulated |
| ENSG00000145817.17 | control-ifn | 6.21E-05 | -0.19149362 | -0.57523528 | down-regulated |
| ENSG00000145901.16 | control-ifn | 4.04E-05 | 0.19764958 | -1.39658255 | down-regulated |
| ENSG00000146063.20 | control-ifn | 0.00604243 | 0.13708563 | 0.317662372 | up-regulated |
| ENSG00000146192.15 | control-ifn | 0.0005738 | 0.3049334 | 0.077617168 | up-regulated |
| ENSG00000146457.16 | control-ifn | 1.73E-05 | -0.11645088 | -0.35713043 | down-regulated |
| ENSG00000146556.15 | control-ifn | 0.00828268 | -0.36081684 | 0.224200266 | up-regulated |
| ENSG00000146828.18 | control-ifn | 1.39E-06 | 0.3164781 | 0.658199059 | up-regulated |
| ENSG00000147050.17 | control-ifn | 0.00046879 | -0.27003504 | -0.05791345 | down-regulated |
| ENSG00000147121.16 | control-ifn | 0.00942716 | 0.18189783 | -0.03291805 | down-regulated |
| ENSG00000147130.14 | control-ifn | 0.00193741 | 0.28313556 | 0.114747959 | up-regulated |
| ENSG00000147416.11 | control-ifn | 9.48E-05 | -0.11957435 | -1.34506166 | down-regulated |
| ENSG00000147548.17 | control-ifn | 3.94E-05 | -0.11061579 | 0.579305954 | up-regulated |
| ENSG00000147854.17 | control-ifn | 0.00473342 | -0.17356749 | 0.573495193 | up-regulated |
| ENSG00000147894.17 | control-ifn | 0.00385757 | 0.37403627 | 0.516898535 | up-regulated |
| ENSG00000148153.14 | control-ifn | 0.00121438 | 0.400568 | -0.74965769 | down-regulated |
| ENSG00000148180.21 | control-ifn | 0.00206515 | 0.17868652 | -0.24410271 | down-regulated |
| ENSG00000148341.18 | control-ifn | 0.00962067 | 0.13606383 | 0.574629186 | up-regulated |
| ENSG00000148396.19 | control-ifn | 6.74E-05 | 0.1423326 | -0.98105939 | down-regulated |
| ENSG00000148688.14 | control-ifn | 0.00133398 | 0.20480692 | -0.0505987 | down-regulated |
| ENSG00000148842.18 | control-ifn | 0.00337424 | 0.345281 | 1.637484912 | up-regulated |
| ENSG00000149177.14 | control-ifn | 0.00847791 | -0.21376736 | -0.77845878 | down-regulated |
| ENSG00000149187.19 | control-ifn | 2.53E-06 | -0.10564583 | -0.50272138 | down-regulated |
| ENSG00000149483.13 | control-ifn | 3.90E-09 | 0.20769723 | -1.310918 | down-regulated |
| ENSG00000149716.13 | control-ifn | 0.00265476 | -0.32956418 | -0.16008098 | down-regulated |
| ENSG00000149761.9 | control-ifn | 0.00956846 | 0.10923481 | -0.01553818 | down-regulated |
| ENSG00000149930.18 | control-ifn | 5.39E-05 | 0.27679362 | -0.63268689 | down-regulated |
| ENSG00000149932.17 | control-ifn | 0.00405916 | 0.11015975 | 0.859768588 | up-regulated |
| ENSG00000150347.17 | control-ifn | 0.00187689 | -0.16982018 | -0.45719795 | down-regulated |
| ENSG00000150593.18 | control-ifn | 1.25E-06 | -0.17028849 | -1.59071939 | down-regulated |
| ENSG00000150637.9 | control-ifn | 0.00095971 | -0.11882658 | -0.34564644 | down-regulated |
| ENSG00000150712.11 | control-ifn | 7.39E-05 | -0.36392212 | 0.119617961 | up-regulated |
| ENSG00000151247.13 | control-ifn | 0.00103311 | 0.14185722 | -0.32619896 | down-regulated |

| | | | | | |
|---|---|---|---|---|---|
| ENSG00000151490.15 | control-ifn | 0.0005738 | -0.30865394 | -1.51674144 | down-regulated |
| ENSG00000151500.15 | control-ifn | 0.00867731 | 0.23996735 | -0.1611816 | down-regulated |
| ENSG00000151576.10 | control-ifn | 0.0043022 | -0.20090359 | -0.2130715 | down-regulated |
| ENSG00000151651.16 | control-ifn | 0.00356006 | -0.10905227 | 0.245890449 | up-regulated |
| ENSG00000151726.15 | control-ifn | 0.00189199 | -0.12503827 | -2.94546321 | down-regulated |
| ENSG00000151883.19 | control-ifn | 0.00340456 | 0.19867243 | -0.00843792 | down-regulated |
| ENSG00000152223.16 | control-ifn | 0.00656426 | -0.25677671 | -0.57588757 | down-regulated |
| ENSG00000152939.17 | control-ifn | 0.00023715 | -1 | -0.60966975 | down-regulated |
| ENSG00000153037.15 | control-ifn | 0.0019847 | 0.14696155 | -0.23027345 | down-regulated |
| ENSG00000153066.13 | control-ifn | 0.0010391 | -0.1430683 | 0.714782749 | up-regulated |
| ENSG00000153107.13 | control-ifn | 0.00368254 | 0.16389726 | 0.053552641 | up-regulated |
| ENSG00000153234.15 | control-ifn | 0.00044013 | -0.2753274 | 4.150286637 | up-regulated |
| ENSG00000153551.14 | control-ifn | 0.00508214 | 0.10550354 | 0.406803259 | up-regulated |
| ENSG00000153786.13 | control-ifn | 0.00536358 | 0.14375138 | 1.023741853 | up-regulated |
| ENSG00000153827.14 | control-ifn | 0.00046533 | 0.10168365 | -1.12334954 | down-regulated |
| ENSG00000153914.16 | control-ifn | 0.00337424 | -0.14987134 | -0.43117702 | down-regulated |
| ENSG00000154370.16 | control-ifn | 0.00095235 | -0.10438112 | 0.169757773 | up-regulated |
| ENSG00000154451.15 | control-ifn | 0.000827 | -0.13670006 | -2.18211744 | down-regulated |
| ENSG00000154781.17 | control-ifn | 0.00680243 | -0.19671231 | -0.0627192 | down-regulated |
| ENSG00000155229.21 | control-ifn | 0.0039559 | -0.20560525 | -0.21672855 | down-regulated |
| ENSG00000155329.12 | control-ifn | 0.0033443 | 0.34478333 | -0.03013393 | down-regulated |
| ENSG00000155363.19 | control-ifn | 0.00062079 | -0.10173378 | -3.25067206 | down-regulated |
| ENSG00000155508.14 | control-ifn | 0.00203984 | -0.13975354 | 0.41558191 | up-regulated |
| ENSG00000155755.20 | control-ifn | 0.00284468 | 0.24041398 | 0.097758861 | up-regulated |
| ENSG00000155926.14 | control-ifn | 1.93E-06 | -0.11779641 | 0.035042173 | up-regulated |
| ENSG00000156256.15 | control-ifn | 0.00528726 | -0.16634494 | -0.14475002 | down-regulated |
| ENSG00000156502.14 | control-ifn | 2.35E-05 | 0.20425958 | 0.448733729 | up-regulated |
| ENSG00000156735.11 | control-ifn | 0.00638233 | -0.30445337 | 0.797407726 | up-regulated |
| ENSG00000156873.16 | control-ifn | 0.00140223 | -0.13312261 | 0.320813315 | up-regulated |
| ENSG00000156976.17 | control-ifn | 8.78E-11 | -0.21116746 | 0.548735173 | up-regulated |
| ENSG00000156990.14 | control-ifn | 0.00014385 | -0.11441657 | 0.090569625 | up-regulated |
| ENSG00000157106.18 | control-ifn | 5.05E-07 | -0.26781094 | -0.56881586 | down-regulated |
| ENSG00000157500.12 | control-ifn | 0.00036669 | -0.24313617 | 0.120039201 | up-regulated |
| ENSG00000157601.15 | control-ifn | 6.36E-06 | -0.27243274 | -5.17813238 | down-regulated |
| ENSG00000157954.15 | control-ifn | 0.00715591 | 0.1298773 | -0.13936875 | down-regulated |
| ENSG00000158019.21 | control-ifn | 0.00046732 | 0.27741038 | 0.9879966 | up-regulated |
| ENSG00000158122.12 | control-ifn | 0.00313723 | 0.31097877 | 0.593733297 | up-regulated |
| ENSG00000158417.11 | control-ifn | 0.00837455 | -0.13474605 | -0.80521986 | down-regulated |
| ENSG00000158711.14 | control-ifn | 0.00010756 | -0.25938042 | -0.61052151 | down-regulated |

| | | | | | |
|---|---|---|---|---|---|
| ENSG00000158805.12 | control-ifn | 0.00011576 | -0.14613751 | -0.33393528 | down-regulated |
| ENSG00000158941.17 | control-ifn | 7.39E-05 | -0.12342613 | -0.73978016 | down-regulated |
| ENSG00000159110.21 | control-ifn | 0.00049315 | -0.15350891 | 0.405571391 | up-regulated |
| ENSG00000159140.22 | control-ifn | 1.41E-08 | 0.18347953 | -0.54493252 | down-regulated |
| ENSG00000159216.19 | control-ifn | 0.00289578 | -0.24043909 | 0.325989855 | up-regulated |
| ENSG00000159322.18 | control-ifn | 0.0015688 | 0.13736119 | -0.16957653 | down-regulated |
| ENSG00000159377.11 | control-ifn | 8.57E-05 | -0.19894909 | 0.220969008 | up-regulated |
| ENSG00000159459.12 | control-ifn | 0.00011576 | -0.24129311 | -0.38455415 | down-regulated |
| ENSG00000159579.14 | control-ifn | 0.0064862 | -0.17380185 | -0.96246807 | down-regulated |
| ENSG00000159658.15 | control-ifn | 0.00037318 | 0.22502937 | -0.5253633 | down-regulated |
| ENSG00000160570.14 | control-ifn | 1.05E-05 | 0.32376139 | 0.611242893 | up-regulated |
| ENSG00000160679.13 | control-ifn | 3.48E-07 | 0.25290945 | -0.59195712 | down-regulated |
| ENSG00000160710.18 | control-ifn | 1.85E-08 | 0.13156286 | -2.14759695 | down-regulated |
| ENSG00000160785.14 | control-ifn | 0.00224323 | -0.14338883 | -0.88962864 | down-regulated |
| ENSG00000160789.24 | control-ifn | 0.00013238 | -0.3695286 | -1.13881169 | down-regulated |
| ENSG00000160799.12 | control-ifn | 0.00349852 | -0.30991383 | 0.835598248 | up-regulated |
| ENSG00000160932.11 | control-ifn | 0.00226487 | 0.11340303 | -2.41874299 | down-regulated |
| ENSG00000160959.8 | control-ifn | 0.00061826 | 0.20216052 | -0.31431029 | down-regulated |
| ENSG00000161217.12 | control-ifn | 0.00201986 | -0.12494319 | -0.93921871 | down-regulated |
| ENSG00000161526.15 | control-ifn | 0.00014189 | -0.14632841 | 0.095639708 | up-regulated |
| ENSG00000161791.14 | control-ifn | 0.00254536 | 0.26067909 | -1.95485722 | down-regulated |
| ENSG00000162174.12 | control-ifn | 0.00201256 | -0.81611325 | 2.540733334 | up-regulated |
| ENSG00000162236.13 | control-ifn | 0.0061495 | -0.10146897 | -0.2687131 | down-regulated |
| ENSG00000162384.14 | control-ifn | 0.00113823 | 0.12950312 | 0.167195244 | up-regulated |
| ENSG00000162402.14 | control-ifn | 0.00189199 | -0.27520577 | -0.36776647 | down-regulated |
| ENSG00000162433.15 | control-ifn | 0.00059834 | 0.43076469 | -3.788855 | down-regulated |
| ENSG00000162650.17 | control-ifn | 0.00824074 | -0.16135816 | -0.21999288 | down-regulated |
| ENSG00000162695.12 | control-ifn | 8.57E-05 | 0.28214032 | -1.25029234 | down-regulated |
| ENSG00000162711.18 | control-ifn | 0.00331446 | -0.25160466 | 0.960755927 | up-regulated |
| ENSG00000162852.14 | control-ifn | 0.00486174 | -0.42775191 | 0.752467184 | up-regulated |
| ENSG00000163110.15 | control-ifn | 2.69E-05 | 0.18751707 | -0.29120503 | down-regulated |
| ENSG00000163156.12 | control-ifn | 0.00022194 | 0.25715706 | 0.743580922 | up-regulated |
| ENSG00000163219.12 | control-ifn | 7.30E-07 | -0.32182682 | -0.55476089 | down-regulated |
| ENSG00000163291.14 | control-ifn | 0.00903182 | -0.23170689 | 0.023641609 | up-regulated |
| ENSG00000163322.14 | control-ifn | 0.00057079 | 0.17012574 | 0.196616594 | up-regulated |
| ENSG00000163399.16 | control-ifn | 0.00034731 | -0.11664726 | -0.22506337 | down-regulated |
| ENSG00000163513.19 | control-ifn | 0.00114915 | 0.12059811 | -0.41212518 | down-regulated |
| ENSG00000163602.12 | control-ifn | 0.00136537 | -0.30649384 | -0.17646648 | down-regulated |
| ENSG00000163644.15 | control-ifn | 0.00149614 | -0.13146482 | -2.48792748 | down-regulated |

| | | | | | |
|---|---|---|---|---|---|
| ENSG00000163684.12 | control-ifn | 0.00865627 | 0.24737932 | 0.635910929 | up-regulated |
| ENSG00000163807.6 | control-ifn | 0.00959021 | 0.19506161 | 0.635738588 | up-regulated |
| ENSG00000163820.16 | control-ifn | 0.0028656 | -0.13180538 | -0.79105608 | down-regulated |
| ENSG00000163872.16 | control-ifn | 0.00189199 | 0.18568286 | -1.56604642 | down-regulated |
| ENSG00000163877.11 | control-ifn | 0.00101289 | -0.13721489 | -0.00256699 | down-regulated |
| ENSG00000163930.10 | control-ifn | 0.00047346 | -0.15008356 | 0.234414565 | up-regulated |
| ENSG00000163939.18 | control-ifn | 6.76E-07 | -0.1391686 | -0.31987239 | down-regulated |
| ENSG00000164077.15 | control-ifn | 0.0005738 | -0.50529953 | -1.23466533 | down-regulated |
| ENSG00000164088.18 | control-ifn | 0.00036867 | 0.13871035 | 0.17646407 | up-regulated |
| ENSG00000164180.14 | control-ifn | 0.00983549 | -0.26424009 | -0.10611145 | down-regulated |
| ENSG00000164190.19 | control-ifn | 0.0018164 | 0.14091585 | -0.60505747 | down-regulated |
| ENSG00000164329.14 | control-ifn | 0.00023579 | -0.16951361 | -0.05607072 | down-regulated |
| ENSG00000164347.18 | control-ifn | 0.00310176 | -0.32689673 | -0.56193674 | down-regulated |
| ENSG00000164463.12 | control-ifn | 3.03E-06 | -0.4176614 | 1.236608276 | up-regulated |
| ENSG00000164483.17 | control-ifn | 8.42E-05 | -0.30424092 | 0.020065044 | up-regulated |
| ENSG00000164506.15 | control-ifn | 6.69E-05 | -0.32764768 | 0.793802815 | up-regulated |
| ENSG00000164576.12 | control-ifn | 0.00090389 | 0.22092673 | 0.06648964 | up-regulated |
| ENSG00000164620.9 | control-ifn | 0.00451093 | -0.36975072 | 0.518294389 | up-regulated |
| ENSG00000164985.15 | control-ifn | 0.00044318 | 0.26601628 | 0.483346056 | up-regulated |
| ENSG00000165219.23 | control-ifn | 2.32E-06 | -0.19155406 | -0.62262591 | down-regulated |
| ENSG00000165280.18 | control-ifn | 0.00413896 | 0.13453176 | -0.86215607 | down-regulated |
| ENSG00000165525.18 | control-ifn | 4.66E-05 | 0.13237731 | -0.49972529 | down-regulated |
| ENSG00000165702.15 | control-ifn | 0.00774385 | 0.61504564 | 1.516557611 | up-regulated |
| ENSG00000165792.18 | control-ifn | 0.00432073 | 0.35930804 | -0.01320855 | down-regulated |
| ENSG00000165813.20 | control-ifn | 0.00256022 | 0.26172755 | -0.06610136 | down-regulated |
| ENSG00000165819.12 | control-ifn | 0.00984948 | -0.18193933 | -0.24913066 | down-regulated |
| ENSG00000165914.15 | control-ifn | 0.00581098 | 0.22926994 | -2.71260892 | down-regulated |
| ENSG00000166037.11 | control-ifn | 0.00522103 | -0.20400203 | 0.095336666 | up-regulated |
| ENSG00000166181.13 | control-ifn | 0.00121432 | -0.11767189 | -0.56056301 | down-regulated |
| ENSG00000166260.13 | control-ifn | 0.00034881 | 0.12530075 | -0.03417479 | down-regulated |
| ENSG00000166266.14 | control-ifn | 0.00430046 | -0.57949418 | 1.209007816 | up-regulated |
| ENSG00000166579.16 | control-ifn | 0.00135792 | -0.26255251 | 0.698258515 | up-regulated |
| ENSG00000166676.17 | control-ifn | 0.00346255 | -0.95322715 | -0.40021024 | down-regulated |
| ENSG00000166797.11 | control-ifn | 0.00280967 | -0.21962823 | 0.430045923 | up-regulated |
| ENSG00000166801.17 | control-ifn | 2.08E-06 | -0.15476033 | -0.23871209 | down-regulated |
| ENSG00000166900.17 | control-ifn | 0.00352647 | 0.34533223 | 0.170366076 | up-regulated |
| ENSG00000166912.17 | control-ifn | 0.00039561 | -0.23163308 | 0.499680692 | up-regulated |
| ENSG00000167005.14 | control-ifn | 0.00201256 | -0.23429641 | 0.620955381 | up-regulated |
| ENSG00000167202.12 | control-ifn | 0.00243555 | -0.40603155 | -0.11327049 | down-regulated |

| | | | | | |
|---|---|---|---|---|---|
| ENSG00000167261.14 | control-ifn | 0.00451093 | 0.17801877 | 1.177613894 | up-regulated |
| ENSG00000167380.17 | control-ifn | 0.00280967 | 0.54399132 | -0.39876212 | down-regulated |
| ENSG00000167461.12 | control-ifn | 0.00842365 | 0.16848558 | -0.45802669 | down-regulated |
| ENSG00000167483.19 | control-ifn | 0.0004373 | -0.33569634 | 2.064542471 | up-regulated |
| ENSG00000167528.13 | control-ifn | 0.0004923 | -0.33082515 | -0.58279946 | down-regulated |
| ENSG00000167637.18 | control-ifn | 0.00033677 | -0.36011458 | -0.44942545 | down-regulated |
| ENSG00000167842.16 | control-ifn | 1.71E-05 | -0.12707183 | -0.00026214 | down-regulated |
| ENSG00000168010.11 | control-ifn | 1.94E-12 | 0.19692245 | 0.653541737 | up-regulated |
| ENSG00000168216.13 | control-ifn | 0.00275338 | 0.17722214 | 0.28110573 | up-regulated |
| ENSG00000168310.12 | control-ifn | 0.00470284 | -0.12023342 | -1.57379728 | down-regulated |
| ENSG00000168395.16 | control-ifn | 0.00025279 | 0.2115159 | 0.421779279 | up-regulated |
| ENSG00000168487.20 | control-ifn | 0.0053075 | 0.23488234 | 0.426061942 | up-regulated |
| ENSG00000168488.19 | control-ifn | 2.26E-06 | -0.10155835 | -0.12519435 | down-regulated |
| ENSG00000168495.13 | control-ifn | 0.00109733 | -0.34258556 | -1.03171154 | down-regulated |
| ENSG00000168675.19 | control-ifn | 0.00271958 | -0.15686784 | -0.21508427 | down-regulated |
| ENSG00000168685.15 | control-ifn | 9.16E-06 | -0.1135741 | -0.73743527 | down-regulated |
| ENSG00000168701.19 | control-ifn | 0.00269481 | 0.1318569 | 0.086384135 | up-regulated |
| ENSG00000168827.15 | control-ifn | 0.00926557 | 0.22397482 | 0.027962145 | up-regulated |
| ENSG00000168876.9 | control-ifn | 0.00965381 | -0.13875215 | 0.421395969 | up-regulated |
| ENSG00000168884.15 | control-ifn | 0.00150947 | -0.15047151 | -0.53332042 | down-regulated |
| ENSG00000168906.13 | control-ifn | 7.20E-08 | -0.36167775 | -1.01581133 | down-regulated |
| ENSG00000169018.6 | control-ifn | 0.0010486 | -0.26538561 | 0.673804715 | up-regulated |
| ENSG00000169062.15 | control-ifn | 0.0091275 | -0.20757502 | 0.063427502 | up-regulated |
| ENSG00000169155.10 | control-ifn | 0.00015172 | 0.41096597 | 0.398166187 | up-regulated |
| ENSG00000169228.14 | control-ifn | 0.00094811 | 0.20374693 | 0.502086614 | up-regulated |
| ENSG00000169239.13 | control-ifn | 0.0060345 | 0.23989409 | 2.718364483 | up-regulated |
| ENSG00000169375.16 | control-ifn | 1.73E-05 | -0.21416053 | -0.25647367 | down-regulated |
| ENSG00000169592.15 | control-ifn | 1.85E-08 | -0.17830094 | 0.995027974 | up-regulated |
| ENSG00000169660.17 | control-ifn | 0.00130987 | -0.16399119 | -0.78701935 | down-regulated |
| ENSG00000169683.8 | control-ifn | 0.00938638 | 0.24457068 | 0.567710875 | up-regulated |
| ENSG00000169756.17 | control-ifn | 0.00010952 | -0.18272619 | -1.28500291 | down-regulated |
| ENSG00000170185.11 | control-ifn | 2.13E-05 | 0.25582769 | 0.23861081 | up-regulated |
| ENSG00000170473.17 | control-ifn | 0.00364159 | -0.28037628 | 0.048200675 | up-regulated |
| ENSG00000170776.22 | control-ifn | 1.53E-05 | -0.12030775 | 0.229495281 | up-regulated |
| ENSG00000171174.15 | control-ifn | 0.00713296 | -0.37436168 | 2.752770878 | up-regulated |
| ENSG00000171302.17 | control-ifn | 0.00237868 | 0.50307726 | -0.74606907 | down-regulated |
| ENSG00000171310.11 | control-ifn | 0.00114915 | -0.24501658 | 1.105975456 | up-regulated |
| ENSG00000171791.14 | control-ifn | 6.83E-08 | -0.2329529 | 1.255805419 | up-regulated |
| ENSG00000171988.20 | control-ifn | 0.00350721 | 0.10339233 | 0.166344871 | up-regulated |

| | | | | | |
|---|---|---|---|---|---|
| ENSG00000172366.20 | control-ifn | 0.00011434 | 0.19494864 | 1.248685096 | up-regulated |
| ENSG00000172375.14 | control-ifn | 0.00140223 | 0.14032229 | 0.298164615 | up-regulated |
| ENSG00000172466.16 | control-ifn | 0.000827 | 0.22553291 | 0.112257165 | up-regulated |
| ENSG00000172575.13 | control-ifn | 0.00037696 | 0.16488817 | -0.51822327 | down-regulated |
| ENSG00000172578.12 | control-ifn | 1.33E-05 | -0.23414572 | -0.72922383 | down-regulated |
| ENSG00000172590.18 | control-ifn | 0.00056528 | 0.23240434 | 0.470216461 | up-regulated |
| ENSG00000172725.14 | control-ifn | 0.00067346 | 0.34716085 | 0.416583968 | up-regulated |
| ENSG00000172795.17 | control-ifn | 2.85E-06 | 0.3662636 | 0.371589565 | up-regulated |
| ENSG00000172845.18 | control-ifn | 0.00074441 | 0.15506539 | 0.452584093 | up-regulated |
| ENSG00000172869.15 | control-ifn | 0.00036669 | -0.50280008 | 0.397697531 | up-regulated |
| ENSG00000172922.11 | control-ifn | 0.00191384 | -0.16510297 | 0.393896517 | up-regulated |
| ENSG00000172932.16 | control-ifn | 2.37E-06 | 0.13797855 | -0.08168774 | down-regulated |
| ENSG00000173153.17 | control-ifn | 0.00642626 | 0.17983359 | -0.13415553 | down-regulated |
| ENSG00000173221.14 | control-ifn | 4.65E-06 | 0.13865925 | -0.29977793 | down-regulated |
| ENSG00000173273.16 | control-ifn | 0.00417752 | -0.47679358 | 0.293835271 | up-regulated |
| ENSG00000173442.13 | control-ifn | 0.00608206 | -0.31463486 | -1.11621192 | down-regulated |
| ENSG00000173451.7 | control-ifn | 9.23E-05 | -0.33344159 | 0.304875005 | up-regulated |
| ENSG00000173575.22 | control-ifn | 0.00029697 | -0.11730871 | -0.16736863 | down-regulated |
| ENSG00000173744.18 | control-ifn | 0.00335071 | 0.14756464 | -1.02424607 | down-regulated |
| ENSG00000173812.11 | control-ifn | 2.02E-08 | 0.16043159 | 1.385276495 | up-regulated |
| ENSG00000173846.13 | control-ifn | 8.88E-05 | -0.25366218 | 1.227727288 | up-regulated |
| ENSG00000173889.16 | control-ifn | 0.00019445 | -0.18004549 | 0.301249647 | up-regulated |
| ENSG00000173933.21 | control-ifn | 0.00984948 | -0.12036383 | -0.03528868 | down-regulated |
| ENSG00000174231.17 | control-ifn | 0.00661954 | -0.18299002 | 0.221284744 | up-regulated |
| ENSG00000174437.18 | control-ifn | 1.09E-05 | -0.12715302 | -0.49214836 | down-regulated |
| ENSG00000174606.14 | control-ifn | 0.00082717 | 0.17215208 | -0.34331195 | down-regulated |
| ENSG00000175087.10 | control-ifn | 0.00726035 | -0.32364913 | -0.14837958 | down-regulated |
| ENSG00000175274.19 | control-ifn | 0.00504083 | 0.19935076 | 0.78241213 | up-regulated |
| ENSG00000175309.15 | control-ifn | 7.42E-07 | 0.13626609 | -0.30848714 | down-regulated |
| ENSG00000175354.20 | control-ifn | 2.70E-09 | 0.31176973 | -0.30821653 | down-regulated |
| ENSG00000175470.20 | control-ifn | 1.23E-10 | -0.33625593 | -0.31288039 | down-regulated |
| ENSG00000175595.16 | control-ifn | 2.36E-05 | -0.44831553 | -0.40611427 | down-regulated |
| ENSG00000175634.15 | control-ifn | 0.0002074 | 0.13891123 | 0.291079358 | up-regulated |
| ENSG00000175662.18 | control-ifn | 0.00042955 | -0.34572522 | 0.056567942 | up-regulated |
| ENSG00000175866.16 | control-ifn | 0.00938638 | 0.25999404 | 0.497762972 | up-regulated |
| ENSG00000176715.17 | control-ifn | 0.00533101 | 0.10363348 | -0.46434961 | down-regulated |
| ENSG00000176986.16 | control-ifn | 0.00928271 | -0.15579167 | -0.89886278 | down-regulated |
| ENSG00000177000.13 | control-ifn | 4.71E-06 | -0.16674813 | 0.398017184 | up-regulated |
| ENSG00000177189.14 | control-ifn | 0.00045626 | 0.14606161 | -0.2379484 | down-regulated |

| | | | | | |
|---|---|---|---|---|---|
| ENSG00000177426.22 | control-ifn | 9.09E-07 | 0.2677403 | -0.28000734 | down-regulated |
| ENSG00000177479.20 | control-ifn | 0.00063605 | 0.11370173 | 0.119130351 | up-regulated |
| ENSG00000177963.15 | control-ifn | 0.00022279 | 0.11475787 | -0.54287688 | down-regulated |
| ENSG00000178104.20 | control-ifn | 9.09E-07 | -0.25171269 | -2.75429635 | down-regulated |
| ENSG00000178127.13 | control-ifn | 0.00281339 | -0.11615796 | -0.22377191 | down-regulated |
| ENSG00000178177.17 | control-ifn | 0.00254536 | -0.23907821 | 0.324338673 | up-regulated |
| ENSG00000178188.15 | control-ifn | 0.00438604 | -0.15387999 | -0.25974542 | down-regulated |
| ENSG00000178209.17 | control-ifn | 8.40E-05 | -0.20403033 | -0.50035245 | down-regulated |
| ENSG00000178573.7 | control-ifn | 0.00605116 | -0.2615961 | -0.31769769 | down-regulated |
| ENSG00000178685.14 | control-ifn | 0.00038935 | 0.11801141 | -2.63951368 | down-regulated |
| ENSG00000178927.18 | control-ifn | 0.00025375 | 0.16733746 | -0.46101915 | down-regulated |
| ENSG00000179094.16 | control-ifn | 0.000456 | -0.18054809 | 3.841130995 | up-regulated |
| ENSG00000179262.10 | control-ifn | 0.00160697 | -0.17730838 | -0.1472224 | down-regulated |
| ENSG00000179335.20 | control-ifn | 1.01E-06 | -0.22111281 | 0.759955306 | up-regulated |
| ENSG00000179454.14 | control-ifn | 0.00010463 | -0.33762018 | 0.081902138 | up-regulated |
| ENSG00000179630.11 | control-ifn | 0.00036157 | 0.48403739 | -5.18066471 | down-regulated |
| ENSG00000179632.10 | control-ifn | 0.00293162 | -0.26639208 | 0.669199855 | up-regulated |
| ENSG00000180228.14 | control-ifn | 0.00082696 | 0.19800016 | -0.08884232 | down-regulated |
| ENSG00000181038.14 | control-ifn | 0.0064479 | 0.18287217 | -0.04486667 | down-regulated |
| ENSG00000181090.21 | control-ifn | 2.58E-06 | 0.17252409 | -0.78882313 | down-regulated |
| ENSG00000181904.9 | control-ifn | 0.00112173 | -0.34386449 | 0.302185314 | up-regulated |
| ENSG00000182196.14 | control-ifn | 0.00053629 | 0.11158399 | 0.970108731 | up-regulated |
| ENSG00000182473.22 | control-ifn | 1.22E-10 | 0.16486082 | -0.01224146 | down-regulated |
| ENSG00000182484.15 | control-ifn | 0.00031007 | -0.11256297 | 0.31716384 | up-regulated |
| ENSG00000182831.12 | control-ifn | 0.00528123 | -0.13540098 | 0.771410064 | up-regulated |
| ENSG00000182934.12 | control-ifn | 0.00046533 | -0.13487876 | -0.32187234 | down-regulated |
| ENSG00000183250.13 | control-ifn | 0.00189269 | 0.16096934 | -1.96818701 | down-regulated |
| ENSG00000183291.18 | control-ifn | 0.00758024 | 0.120144 | 0.503051928 | up-regulated |
| ENSG00000183486.14 | control-ifn | 3.62E-15 | -0.11270902 | -4.00722085 | down-regulated |
| ENSG00000183506.17 | control-ifn | 0.00131992 | -0.19457401 | 0.438442003 | up-regulated |
| ENSG00000183696.14 | control-ifn | 6.01E-06 | 0.14952246 | -0.17004345 | down-regulated |
| ENSG00000183785.16 | control-ifn | 0.00815156 | -0.81271582 | 0.976975992 | up-regulated |
| ENSG00000184014.9 | control-ifn | 0.00244741 | 0.23098943 | -0.29779024 | down-regulated |
| ENSG00000184047.20 | control-ifn | 0.00590349 | -0.26324599 | -0.46977781 | down-regulated |
| ENSG00000184203.8 | control-ifn | 0.0020108 | 0.21252722 | 0.13735727 | up-regulated |
| ENSG00000184205.14 | control-ifn | 0.00038935 | 0.12883527 | -0.1454566 | down-regulated |
| ENSG00000184368.16 | control-ifn | 0.00081572 | 1 | -0.83670499 | down-regulated |
| ENSG00000184432.11 | control-ifn | 0.00092588 | -0.14273772 | -1.58400329 | down-regulated |
| ENSG00000184574.10 | control-ifn | 0.00575899 | -0.26855745 | -0.66969974 | down-regulated |

| | | | | | |
|---|---|---|---|---|---|
| ENSG00000184584.13 | control-ifn | 0.00096794 | 0.10378886 | 0.86898074 | up-regulated |
| ENSG00000184634.17 | control-ifn | 6.01E-06 | -0.15594065 | -1.58177636 | down-regulated |
| ENSG00000184787.19 | control-ifn | 0.00069437 | 0.21998908 | 0.357470886 | up-regulated |
| ENSG00000184863.11 | control-ifn | 0.00033432 | -0.2362628 | 1.052563981 | up-regulated |
| ENSG00000185129.8 | control-ifn | 0.00020359 | 0.31392904 | -0.93305251 | down-regulated |
| ENSG00000185324.22 | control-ifn | 6.70E-05 | 0.12966014 | -0.07484072 | down-regulated |
| ENSG00000185504.17 | control-ifn | 0.00430212 | 0.18013322 | -0.00086199 | down-regulated |
| ENSG00000185507.21 | control-ifn | 0.00034158 | -0.26191002 | -3.35844092 | down-regulated |
| ENSG00000185658.14 | control-ifn | 0.0008105 | 0.27378677 | 0.364035437 | up-regulated |
| ENSG00000185963.14 | control-ifn | 2.35E-05 | -0.36656302 | 0.233321053 | up-regulated |
| ENSG00000186174.13 | control-ifn | 0.0051856 | 0.13558471 | -0.33599268 | down-regulated |
| ENSG00000186416.18 | control-ifn | 0.0057877 | -0.2552869 | 0.033027912 | up-regulated |
| ENSG00000186432.10 | control-ifn | 1.29E-05 | -0.10324377 | -0.34625259 | down-regulated |
| ENSG00000186635.15 | control-ifn | 0.00126713 | -0.14639457 | -0.26650469 | down-regulated |
| ENSG00000186827.11 | control-ifn | 0.00414983 | 0.2050719 | -2.09740131 | down-regulated |
| ENSG00000186951.17 | control-ifn | 0.00949641 | -0.17426861 | 0.537352591 | up-regulated |
| ENSG00000187118.14 | control-ifn | 0.00091348 | -0.17485152 | 0.952107404 | up-regulated |
| ENSG00000187239.17 | control-ifn | 0.00849358 | -0.1133021 | 0.065884394 | up-regulated |
| ENSG00000187608.10 | control-ifn | 0.00087371 | 0.11873215 | -5.52047598 | down-regulated |
| ENSG00000188186.11 | control-ifn | 0.00022091 | 0.18009857 | 1.133639641 | up-regulated |
| ENSG00000188313.13 | control-ifn | 9.41E-10 | 0.24861299 | -2.67493778 | down-regulated |
| ENSG00000188603.22 | control-ifn | 0.00136537 | -0.18396636 | 0.134532594 | up-regulated |
| ENSG00000188647.13 | control-ifn | 0.00036452 | 0.28051369 | 0.491644706 | up-regulated |
| ENSG00000189159.16 | control-ifn | 0.00034731 | 0.20457692 | 0.81460066 | up-regulated |
| ENSG00000196247.13 | control-ifn | 5.29E-05 | 0.24128176 | -0.21729466 | down-regulated |
| ENSG00000196267.14 | control-ifn | 0.00066621 | 0.45381927 | 0.225422517 | up-regulated |
| ENSG00000196405.13 | control-ifn | 6.72E-06 | 0.13529397 | -0.59403121 | down-regulated |
| ENSG00000196502.13 | control-ifn | 0.00077001 | 0.28669053 | 3.200981905 | up-regulated |
| ENSG00000196505.11 | control-ifn | 0.00079739 | 0.18771046 | 0.152765231 | up-regulated |
| ENSG00000196535.19 | control-ifn | 0.00189199 | 0.13500729 | 0.798773878 | up-regulated |
| ENSG00000196652.12 | control-ifn | 0.00201256 | -0.34205326 | -0.34434926 | down-regulated |
| ENSG00000196663.16 | control-ifn | 0.00100153 | 0.28206419 | -0.40425569 | down-regulated |
| ENSG00000196670.14 | control-ifn | 0.00625059 | -0.26824937 | -0.29657548 | down-regulated |
| ENSG00000196776.17 | control-ifn | 0.00370118 | -0.10954358 | -0.31404769 | down-regulated |
| ENSG00000196792.12 | control-ifn | 0.0060345 | -0.38450226 | -0.33542176 | down-regulated |
| ENSG00000196839.14 | control-ifn | 0.00849358 | -0.42264787 | -2.65398311 | down-regulated |
| ENSG00000196843.17 | control-ifn | 2.67E-07 | -0.360528 | -0.23048177 | down-regulated |
| ENSG00000197119.13 | control-ifn | 0.00054088 | -0.19611006 | 1.507132739 | up-regulated |
| ENSG00000197170.10 | control-ifn | 0.00330019 | 0.22037058 | 0.183386527 | up-regulated |

| | | | | | |
|---|---|---|---|---|---|
| ENSG00000197321.16 | control-ifn | 0.00020491 | -0.1158161 | 0.649823865 | up-regulated |
| ENSG00000197386.14 | control-ifn | 0.00126713 | -0.14168083 | -0.82742235 | down-regulated |
| ENSG00000197448.14 | control-ifn | 0.00028482 | 0.27323119 | 1.149522861 | up-regulated |
| ENSG00000197536.11 | control-ifn | 0.00195285 | 0.14089872 | -1.47030006 | down-regulated |
| ENSG00000197619.15 | control-ifn | 0.00625023 | -0.37544027 | -0.64382264 | down-regulated |
| ENSG00000197620.11 | control-ifn | 0.00037696 | -0.19521339 | 0.508036323 | up-regulated |
| ENSG00000197841.15 | control-ifn | 0.00079756 | -0.28546897 | -1.15615372 | down-regulated |
| ENSG00000197857.15 | control-ifn | 0.00237474 | 0.18085249 | 0.127427989 | up-regulated |
| ENSG00000197943.11 | control-ifn | 0.00578183 | 0.13719048 | -0.35079802 | down-regulated |
| ENSG00000197971.16 | control-ifn | 0.00035873 | -0.149302 | -0.27239447 | down-regulated |
| ENSG00000198000.12 | control-ifn | 0.00621978 | -0.10252255 | -0.65868464 | down-regulated |
| ENSG00000198001.16 | control-ifn | 0.00151951 | 0.10262715 | -0.21365624 | down-regulated |
| ENSG00000198055.11 | control-ifn | 0.00239698 | 0.10402694 | 1.458137522 | up-regulated |
| ENSG00000198301.12 | control-ifn | 0.00086573 | -0.11746033 | -0.78332946 | down-regulated |
| ENSG00000198373.13 | control-ifn | 0.00047588 | 0.18398718 | 0.284456494 | up-regulated |
| ENSG00000198408.14 | control-ifn | 0.00053629 | -0.1070297 | -0.16946936 | down-regulated |
| ENSG00000198521.12 | control-ifn | 0.00158492 | -0.13705968 | -0.59919267 | down-regulated |
| ENSG00000198589.14 | control-ifn | 0.00405916 | -0.18618228 | -0.57532379 | down-regulated |
| ENSG00000198604.11 | control-ifn | 0.00261512 | 0.19704064 | -1.33252548 | down-regulated |
| ENSG00000198646.14 | control-ifn | 0.00600899 | -0.11107207 | 0.076613683 | up-regulated |
| ENSG00000198689.13 | control-ifn | 0.00125795 | -0.47437898 | -0.7108397 | down-regulated |
| ENSG00000198771.11 | control-ifn | 0.00034731 | -0.13813604 | 1.243266195 | up-regulated |
| ENSG00000198799.12 | control-ifn | 0.00110937 | -0.27804699 | -0.02974496 | down-regulated |
| ENSG00000198900.7 | control-ifn | 3.59E-05 | -0.16111812 | -1.49428758 | down-regulated |
| ENSG00000203896.10 | control-ifn | 4.16E-05 | -0.10291559 | 2.361900428 | up-regulated |
| ENSG00000204152.14 | control-ifn | 0.00412828 | 0.35294144 | 0.756048496 | up-regulated |
| ENSG00000204304.12 | control-ifn | 9.48E-05 | 0.17223799 | 0.163995855 | up-regulated |
| ENSG00000204463.14 | control-ifn | 0.00926557 | -0.10710489 | -0.35000163 | down-regulated |
| ENSG00000204977.10 | control-ifn | 1.21E-05 | -0.19687425 | 0.010156791 | up-regulated |
| ENSG00000205571.14 | control-ifn | 0.00055984 | -0.43404663 | 0.612557425 | up-regulated |
| ENSG00000205744.10 | control-ifn | 0.00061416 | 0.15084551 | 1.194506584 | up-regulated |
| ENSG00000205758.12 | control-ifn | 0.00386795 | 0.13429529 | 0.314720432 | up-regulated |
| ENSG00000205937.12 | control-ifn | 0.00011576 | 0.25587681 | 0.042573216 | up-regulated |
| ENSG00000211460.12 | control-ifn | 1.03E-05 | -0.25165609 | 0.52655902 | up-regulated |
| ENSG00000213015.9 | control-ifn | 0.00040358 | 0.26776934 | 0.227975671 | up-regulated |
| ENSG00000213658.13 | control-ifn | 0.00025375 | 0.11511558 | -0.36803895 | down-regulated |
| ENSG00000213782.7 | control-ifn | 0.00030058 | -0.38686039 | 0.238910862 | up-regulated |
| ENSG00000213923.13 | control-ifn | 0.00604067 | -0.17772746 | 0.206924599 | up-regulated |
| ENSG00000213930.12 | control-ifn | 0.00497911 | -0.12278204 | 0.061510809 | up-regulated |

| | | | | | |
|---|---|---|---|---|---|
| ENSG00000214022.12 | control-ifn | 1.09E-05 | -0.13877411 | 0.361756529 | up-regulated |
| ENSG00000214253.9 | control-ifn | 0.00956846 | 0.12871129 | 0.83303797 | up-regulated |
| ENSG00000214413.9 | control-ifn | 0.0075394 | 0.20280745 | -0.03449322 | down-regulated |
| ENSG00000214941.8 | control-ifn | 0.00194088 | -0.20745019 | 0.999095513 | up-regulated |
| ENSG00000215068.9 | control-ifn | 0.00383369 | 0.13670162 | -0.33261788 | down-regulated |
| ENSG00000215845.11 | control-ifn | 0.0010406 | 0.14991507 | 0.696795894 | up-regulated |
| ENSG00000217555.13 | control-ifn | 0.0063476 | 0.19762268 | 1.585659733 | up-regulated |
| ENSG00000221909.3 | control-ifn | 0.00969765 | -0.42084821 | -2.20750363 | down-regulated |
| ENSG00000221926.13 | control-ifn | 0.00248549 | -0.47642763 | -0.81002506 | down-regulated |
| ENSG00000223768.3 | control-ifn | 0.0051856 | 0.42946649 | -0.00368596 | down-regulated |
| ENSG00000224383.8 | control-ifn | 0.00667881 | -0.22643426 | 0.500558365 | up-regulated |
| ENSG00000225648.6 | control-ifn | 0.00081081 | -0.20283524 | 1.54973009 | up-regulated |
| ENSG00000225830.16 | control-ifn | 0.00140223 | -0.13640343 | -0.07433254 | down-regulated |
| ENSG00000227627.4 | control-ifn | 0.00943381 | 0.40476541 | 1.908793936 | up-regulated |
| ENSG00000229180.8 | control-ifn | 0.00201242 | 0.19512404 | 0.380670761 | up-regulated |
| ENSG00000232119.8 | control-ifn | 0.00030228 | 0.11996686 | 0.306985602 | up-regulated |
| ENSG00000233016.8 | control-ifn | 0.0001341 | 0.12950946 | 1.243436516 | up-regulated |
| ENSG00000235437.9 | control-ifn | 0.0001324 | 0.32691557 | 1.213563857 | up-regulated |
| ENSG00000239713.9 | control-ifn | 0.00049838 | -0.125542 | -1.51910695 | down-regulated |
| ENSG00000239900.14 | control-ifn | 0.00384507 | 0.19484857 | 0.590786011 | up-regulated |
| ENSG00000245164.9 | control-ifn | 3.95E-06 | 0.35060554 | -0.12582622 | down-regulated |
| ENSG00000245552.8 | control-ifn | 0.00398163 | 0.36545716 | -0.39510963 | down-regulated |
| ENSG00000249859.13 | control-ifn | 0.00177557 | 0.30883576 | -0.09101308 | down-regulated |
| ENSG00000255529.9 | control-ifn | 0.00156831 | 0.13846221 | 0.095358406 | up-regulated |
| ENSG00000256594.9 | control-ifn | 0.00147538 | 0.23398572 | 0.30521402 | up-regulated |
| ENSG00000257103.9 | control-ifn | 0.00050866 | -0.21757437 | 0.578385147 | up-regulated |
| ENSG00000257242.9 | control-ifn | 0.00501333 | 0.81965525 | 2.115556287 | up-regulated |
| ENSG00000257621.9 | control-ifn | 2.47E-05 | 0.10646685 | -0.02695647 | down-regulated |
| ENSG00000260078.4 | control-ifn | 0.0012538 | 0.24779543 | 3.504986575 | up-regulated |
| ENSG00000264343.7 | control-ifn | 1.53E-05 | -0.26909908 | 0.658085475 | up-regulated |
| ENSG00000266173.7 | control-ifn | 0.0064862 | -0.17103931 | -0.55589 | down-regulated |
| ENSG00000267121.6 | control-ifn | 3.77E-07 | -0.1957852 | -0.62339599 | down-regulated |
| ENSG00000269609.6 | control-ifn | 0.00030945 | 0.28476387 | -0.04667208 | down-regulated |
| ENSG00000269743.3 | control-ifn | 0.00237229 | -0.15632967 | -0.61712637 | down-regulated |
| ENSG00000272888.8 | control-ifn | 6.28E-07 | 0.10567423 | 1.733681843 | up-regulated |
| ENSG00000273899.5 | control-ifn | 0.00173725 | 0.10630987 | 0.550277002 | up-regulated |
| ENSG00000274265.5 | control-ifn | 0.00927513 | 0.37233998 | -0.57050252 | down-regulated |
| ENSG00000275023.5 | control-ifn | 1.08E-07 | 0.25222621 | -0.82905159 | down-regulated |
| ENSG00000275052.5 | control-ifn | 0.00144601 | 0.22308912 | -0.85580788 | down-regulated |

| | | | | | |
|---|---|---|---|---|---|
| ENSG00000275111.5 | control-ifn | 0.00413167 | 0.63453706 | -3.0083149 | down-regulated |
| ENSG00000277147.7 | control-ifn | 0.00613653 | 0.11389243 | -0.34742931 | down-regulated |
| ENSG00000278053.5 | control-ifn | 0.00122024 | 0.22221896 | -1.08860438 | down-regulated |
| ENSG00000282851.2 | control-ifn | 8.25E-05 | 0.19575382 | -1.86143414 | down-regulated |
| ENSG00000284024.4 | control-ifn | 0.0001599 | 0.40897293 | 0.68810166 | up-regulated |
| ENSG00000284194.3 | control-ifn | 0.00552192 | 0.29036285 | 0.10724134 | up-regulated |

**RT-qPCR primers**

| | |
|---|---|
| CLK1 mRNA FW | CATCGTCGTTCACATGGGAAG |
| CLK1 mRNA REV | CTTCACCTAAAGTATCAACAATTTCATATC |
| CLK1 sk ex4 FW | CGTCGTTCACATGGGATGAAATTG |
| CLK1 sk ex4 REV | GCTACATGTCTACCTCCCGC |
| CLK1 int ret FW | GAAGTGGAAGAAGTAGTTATAAAAGC |
| CLK1 int ret REV | TGTTCCACATGGGATATAAAATTTCC |
| ISG15 FW | TCCTGGTGAGGAATAACAAGGG |
| ISG15 REV | GTCAGCCAGAACAGGTCGTC |
| IFIT1 FW | GAAATATGAATGAAGCCCTGGA |
| IFIT1 REV | GACCTTGTCTCACAGAGTTCTCAA |
| DENV FW | TTGAGTAAACYRTGCTGCCTGTAGCTC |
| DENV REV | GGGTCTCCTCTAACCTCTAGTCCT |
| U6 FW | CTCGCTTCGGCAGCACA |
| U6 REV | AACGCTTCACGAATTTGCGT |

**sgRNAS**

| | |
|---|---|
| FW CLK1 sgRNA1 | **CACC** GGT CGC GCA CCA ACA ACA AAA |
| REV CLK1 sgRNA1 | **AAAC** TTT TGT TGT TGG TGC GCG ACC |
| FW CLK1 sgRNA2 | **CACC** GAA AGC TTG TAA CGC AAT CAC |
| REV CLK1 sgRNA2 | **AAAC** GTG ATT GCG TTA CAA GCT TTC |
| FW CLK1 sgRNA3 | **CACC** GCA GGA CGG TAA GAC GCA GGA |
| REV CLK1 sgRNA3 | **AAAC** TCC TGC GTC TTA CCG TCC TGC |
| FW CLK1 sgRNA4 | **CACC** GAG CAG GCA GTC GTC GCG CCG |
| REV CLK1 sgRNA4 | **AAAC** CGG CGC GAC GAC TGC CTG CTC |
| FW CLK1 sgRNA5 | **CACC** GTT CAC TCC CAG GCT GCA CAG |
| REV CLK1 sgRNA5 | **AAAC** CTG TGC AGC CTG GGA GTG AAC |

**Cloning primers**

| | |
|---|---|
| FW CLK1 BamHI | atggatccatgtacccctacgacgtgcccgactacgccAGACATTCAAAGAGAACTTACTG |
| REV CLK1 EcoRI | tagaattcTCACGTATGCTTTTTAAGTGGG |
| FW ISRE EcoRI | cgagaattcgatccctgccacgctatggag |
| REV ISRE | CCTCGCCCTTGCTCACCATTTCCATGGTGGCTTTACC |
| FW GFP | ATGGTGAGCAAGGGCGAGGAGC |
| REV GFP BamHI | agcggatccCTTGTACAGCTCGTCCATGCCG |
| FW IntISG15 BamHI | ATCGGATCCGTAAGGCAGATGTCACAGGTGG |
| REV IntISG15 BamHI | AGCGGATCCCTGCAGGCGTCACACAGG |
